# Deciphering novel TCF4-driven mechanisms underlying a common triplet repeat expansion-mediated disease

**DOI:** 10.1101/2023.03.29.534731

**Authors:** Nihar Bhattacharyya, Niuzheng Chai, Nathaniel J Hafford-Tear, Amanda N Sadan, Anita Szabo, Christina Zarouchlioti, Jana Jedlickova, Szi Kay Leung, Tianyi Liao, Lubica Dudakova, Pavlina Skalicka, Mohit Parekh, Ismail Moghul, Aaron R Jeffries, Michael E Cheetham, Kirithika Muthusamy, Alison J Hardcastle, Nikolas Pontikos, Petra Liskova, Stephen J Tuft, Alice E Davidson

## Abstract

Fuchs endothelial corneal dystrophy (FECD) is an age-related cause of vision loss, and the most common repeat expansion-mediated disease in humans characterised to date. Up to 80% of European FECD cases have been attributed to expansion of a non-coding CTG repeat element (termed CTG18.1) located within the ubiquitously expressed transcription factor encoding gene, *TCF4*. The non-coding nature of the repeat and the transcriptomic complexity of *TCF4* have made it extremely challenging to experimentally decipher the molecular mechanisms underlying this disease. Here we comprehensively describe CTG18.1 expansion-driven molecular components of disease within primary patient-derived corneal endothelial cells (CECs), generated from a large cohort of individuals with CTG18.1-expanded (Exp+) and CTG 18.1-independent (Exp-) FECD. We employ long-read, short-read, and spatial transcriptomic techniques to interrogate expansion-specific transcriptomic biomarkers. Interrogation of long-read sequencing and alternative splicing analysis of short-read transcriptomic data together reveals the global extent of altered splicing occurring within Exp+ FECD, and unique transcripts associated with CTG18.1-expansions. Similarly, differential gene expression analysis highlights the total transcriptomic consequences of Exp+ FECD within CECs. Furthermore, differential exon usage, pathway enrichment and spatial transcriptomics reveal *TCF4* isoform ratio skewing solely in Exp+ FECD with potential downstream functional consequences. Lastly, exome data from 134 Exp- FECD cases identified rare (minor allele frequency <0.005) and potentially deleterious (CADD>15) *TCF4* variants in 7/134 FECD Exp- cases, suggesting that *TCF4* variants independent of CTG18.1 may increase FECD risk. In summary, our study supports the hypothesis that at least two distinct pathogenic mechanisms, RNA toxicity and *TCF4* isoform-specific dysregulation, both underpin the pathophysiology of FECD. We anticipate these data will inform and guide the development of translational interventions for this common triplet-repeat mediated disease.

**Author’s summary:** Fuchs endothelial corneal dystrophy (FECD) leads to vision loss and is the most common repeat expansion-mediated disease characterised to date; most individuals with FECD harbour a non-coding CTG repeat expansion within the gene *TCF4*. FECD represents an important paradigm for other rare and devastating neurological repeat-mediated diseases, given its relatively mild and tissue-specific nature. Intriguingly, despite *TCF4* being ubiquitously expressed, individuals with FECD only experience corneal disease, and the biological reason for this tissue-specificity remains elusive. Here, we use tissue from 31 individuals with FECD to perform complementary long-read, short-read and spatial transcriptomic analyses to enhance our understanding of mechanisms underpinning this disease. These data highlight that at least two mechanisms, RNA toxicity and *TCF4* isoform dysregulation, underlie the disease state in affected corneal cells. Furthermore, *TCF4* isoform skewing, with evidence of downregulation, suggests this mechanism in part may explain the unique vulnerability of the cornea. In addition, 7/134 FECD expansion negative cases were identified to harbour rare and potentially deleterious *TCF4* variants, further supporting the hypothesis that dysregulation of TCF4 may be key to FECD pathophysiology. Biological insights presented here will guide the development of personalised FECD therapies and may inform the development of repeat-expansion mediated therapies more broadly.

## Introduction

Fuchs endothelial corneal dystrophy (FECD) is a common, age-related, and visually disabling disease that represents a leading indication for corneal transplantation in high-income countries) (1,2). It primarily affects the corneal endothelium, a monolayer of post-mitotic cells on the posterior corneal surface that function to maintain relative corneal dehydration. As the disease progresses, collagenous excrescences (guttae) form on the thickened corneal endothelial basement membrane (Descemet membrane) and corneal endothelial cells (CECs) undergo accelerated cell death, which results in corneal oedema and opacity (3). Affected individuals typically experience symptoms in their 5^th^ to 6^th^ decade, with glare, reduced contrast sensitivity, and blurred vision (4). Transplantation of donor endothelial tissue is the preferred treatment for cases with visually significant symptoms. However, given an increasing societal burden of age-related disease coupled with a global shortage of suitable donor tissue, there is a pressing clinical need for alternative treatments (3,5).

Up to 80% of FECD cases of European descent have one or more expanded copies of a CTG trinucleotide repeat element (termed CTG18.1; MIM #613267) located within an intronic region of the transcription factor gene 4, *TCF4* (4,6). We have previously demonstrated, in our large (n=450) FECD cohort, that an expanded copy of this CTG repeat (defined as ≥50 copies) confers >76-fold increased risk for developing FECD (7). Expansion of non-coding repeat elements within the human genome have been identified to cause more than 30 human diseases to date (8), including fragile X tremor ataxia syndrome (MIM #300623), myotonic dystrophy types 1 and 2 (MIM #160900 and #602668) and *C9orf72*-associated amyotrophic lateral sclerosis (ALS) and frontotemporal dementia (MIM #105550)(8). Expansion-positive (Exp+) FECD represents by far the most common disease within this category. However, to date, it has only been established that it affects the cornea, which is in contrast to the majority of the other much rarer non-coding repeat expansion disorders that typically have devastating outcomes on neurological and/or neuromuscular systems. Despite these differences, strong mechanistic molecular parallels exist between CTG18.1-mediated FECD and other non-coding repeat expansion diseases, whereby multiple pathogenic mechanisms occur concurrently to drive pathology (8,9).

We, and others, have demonstrated that RNA toxicity comprising splicing dysregulation attributed to the accumulation of CUG-containing RNA foci and sequestration of splicing factors MBNL1 and MBNL2 within affected cells, is a key feature of FECD in the presence of one or more expanded CTG18.1 alleles (6,7,10–12). Alongside this RNA induced toxicity, repeat-associated non-AUG (RAN) dependent translation of the expanded repeat, and dysregulation of *TCF4* itself have also been hypothesised to contribute to the underlying pathology of CTG18.1 Exp+ mediated FECD(13). However, a comprehensive understanding of consequences and role of both the spliceopathy and downstream effects of the *TCF4* dysregulation have yet to be characterised in parallel in the same model system. Furthermore, because *TCF4* comprises >90 alternative spatially and temporally expressed isoforms, deciphering transcript-specific level dysregulation has proven to be extremely challenging and a consistent pattern of *TCF4* dysregulation has yet to be associated with CTG18.1 expansions in corneal endothelial cells (CECs) (14–16). Such diversity is reflected in the varied functional roles TCF4 plays in human development, alongside the range of human disease associated with *TCF4* variants in addition to FECD. For example, *TCF4* haploinsufficiency results in Pitt-Hopkins syndrome (MIM #610954) (17,18), a severe neurodevelopmental disorder, whereas somatic *TCF4* mutations are commonly identified in Sonic hedgehog medulloblastoma (19) and common germline variants have been associated with Schizophrenia risk (20,21).

In this study, we used a large cohort of biologically independent primary FECD case-derived CECs to isolate the effects of the multiple parallel pathogenic processes underlying CTG18.1-mediated FECD. We present long-read transcriptomic data to enhance our understanding of the increased isoform diversity observed in Exp+ FECD, and spatial transcriptomics to demonstrate *TCF4*-isoform dysregulation. For the first time, we also isolate downstream transcriptomic signatures of dysregulation that we hypothesise to be driven by *TCF4-*isoform dysregulation that may, in part, explain the tissue specificity of the disease. We go on to explore the potential relevance of TCF4 dysregulation to FECD, independent of the CTG18.1 expansion, by interrogating expansion-negative FECD cases for rare *TCF4* variants through exome sequencing and subsequently applying a gene-burden style approach.

## Results

### RNA-seq data from CECs reveal CTG18.1 expansion-mediated FECD to be transcriptomically distinct from non-expanded FECD

To further increase our understanding of CTG18.1 expansion-associated splicing dysregulation events, and better understand the relationship between individual events to transcript-specific isoforms, we generated long-read RNA-seq data from biologically independent unpassaged primary CEC cultures derived from healthy control (n=4, mean age 68.75 years) and Exp+ FECD (mono-allelic CTG18.1 allele repeat length ≥50) (n=4, mean age 66.8 years) tissues (**Table 1, S1**). After alignment and stringent quality control, reads that displayed evidence of 5’ degradation, intrapriming and reverse transcription template switching were removed. Exp+ and control samples were subsequently analysed with SQANTI3 to assess isoform diversity (**Figure 1A, B and Table 2, S2**). We also generated short-read RNA-seq data from unpassaged primary CEC cultures derived from healthy control (n=4, mean age 58.5 years), Exp+ FECD (mono-allelic CTG18.1 allele repeat length ≥50) (n=3, mean age 63.3 years) and Exp- FECD (bi-allelic CTG18.1 allele repeat lengths ≤30) (n=3, mean age 63.3 years) tissues (**Table 1**). Exome data generated from the Exp- FECD cases utilised in this transcriptomic work were analysed for the presence of variants in any previously FECD-associated genes, such as *TCF4, COL8A2, SLC4A11, ZEB1,* and *AGBL1*(4), to identify any potential underlying genetic cause. No rare (MAF <0.005) or potentially deleterious (CADD >15) variants were identified. Hierarchical clustering using heatmap visualisation and PCA analysis including sex and ethnicity as covariates revealed that Exp+ samples grouped separately from not only the controls, but also the Exp- samples **(Figure 1C, D)**. Driving genes and additional principal components are listed in Supplemental Table S2. Notably, Exp- samples also clustered together on the PCA plot, but the heatmap derived from hierarchical clustering highlighted that greater intra-group variability existed between Exp- samples compared to variability within control and Exp+ groups (**Figure 1C**). We hypothesise that the increased transcriptomic diversity within the Exp- group is due to potentially distinct, as yet unidentified, genetic causes driving disease for each FECD case-derived sample. We included all three groups in the downstream comparative pipelines using the following pairwise comparisons; **(PWC1)** control versus Exp+ FECD, **(PWC2)** Exp+ FECD versus Exp- FECD, and **(PWC3)** control versus Exp- FECD.

**Figure 1:**
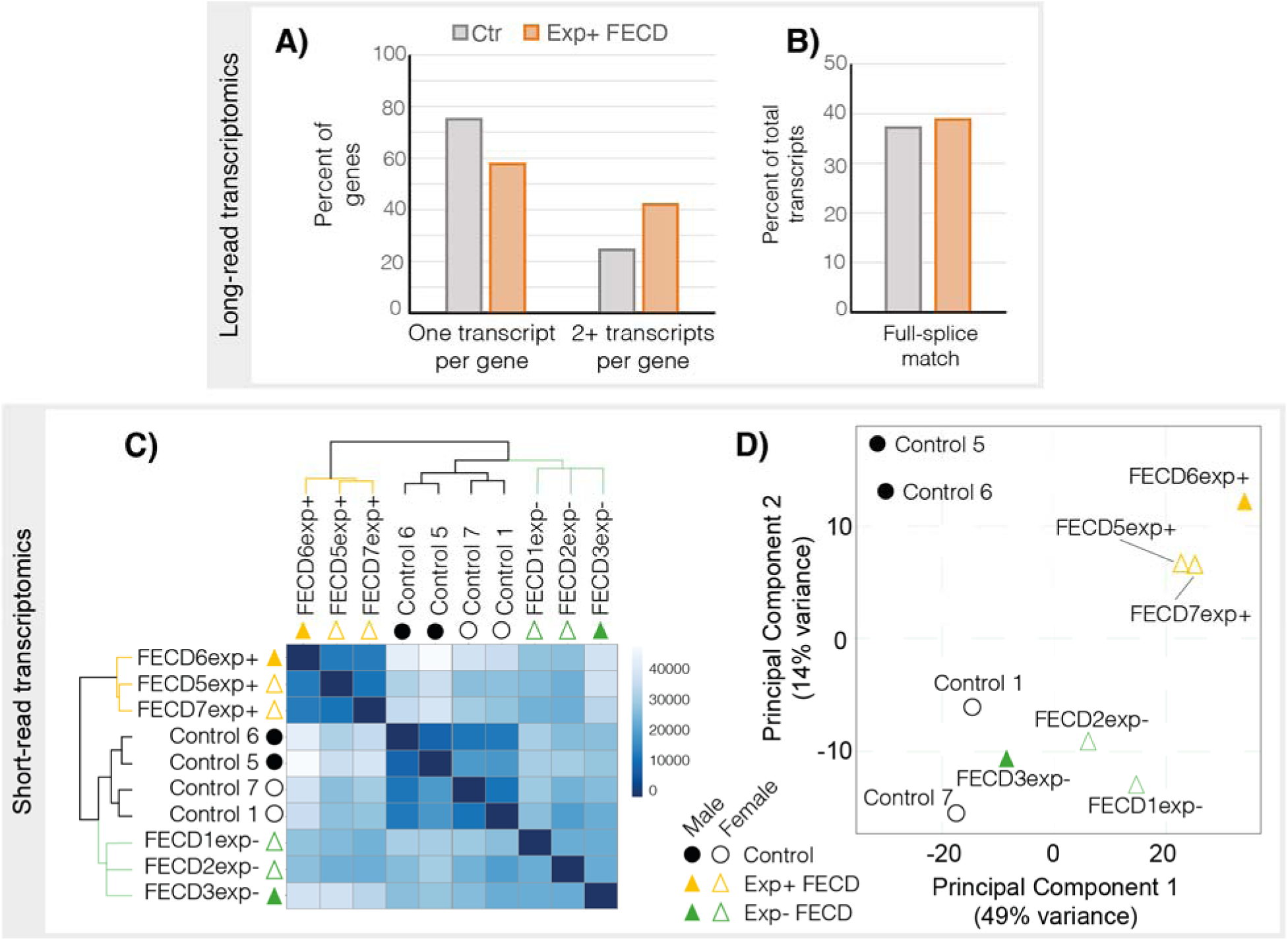
Exp+ FECD samples are transcriptomically distinct in both short- and long-read datasets when compared to controls. **A)** Bar chart showing proportion of annotated genes in long read transcriptomic dataset with only one transcript detect vs 2 or more transcripts in Control and Exp+ samples. Exp+ long-read samples have a higher proportion of genes with more than one isoform identified. **B)** Proportion of annotated transcripts with full splice matches. Exp+ FECD and Control samples have similar percentages of full splice matches suggesting additional splicing diversity seen in Exp+ is not predominantly due to aberrant splicing. **C)** Hierarchical clustering heatmap of sample-to-sample distances within short-read RNA-Seq data. Darker colours denote higher similarities between samples, demonstrating that Exp+ and control are more similar within each subgroup than with other samples. Exp- shows a similar result but is less pronounced, likely due to differences in unidentified underlying genetic causes. **D)** Principal component plot (PCA) of all short-read samples with sex, disease state and ethnicity as covariates. All three subsets of data cluster separately in the PCA plot. Control samples appear as circles, FECD Exp+ appear as orange triangles, FECD Exp- appear as green triangles. Male individuals are represented as filled shapes while female individuals are outlined.

**Table 1:** Summary of FECD case and control-derived corneal endothelial cell (CEC) cultures analysed by short and long-read RNA-seq, and spatial transcriptomics.

| Sample | Sex | Age* | Ethnicity<sup>a</sup>/Ancestry<sup>b</sup> |
| --- | --- | --- | --- |
| *Long-read* |  |  |  |
| Control 1 | Female | 68 | White-European<sup>a</sup> |
| Control 2 | Male | 64 | White-European<sup>a</sup> |
| Control 3 | Female | 74 | White-European<sup>a</sup> |
| Control 4 | Male | 69 | White-European<sup>a</sup> |
| FECD1exp+ (43/90) | Female | 74 | White-European<sup>a,b</sup> |
| FECD2exp+ (14/80) | Male | 78 | White-European<sup>a,b</sup> |
| FECD3exp+ (18/86) | Female | 70 | White-European<sup>a,b</sup> |
| FECD4exp+ (26/78) | Female | 45 | White-European<sup>b</sup> |
| *Short-read* |  |  |  |
| Control 1 | Female | 68 | White-European<sup>a</sup> |
| Control 5 | Male | 55 | White-European<sup>a</sup> |
| Control 6 |  |  |  |
| Control 7 | Female | 56 | Black-African<sup>a</sup> |
| FECD5exp+ (16/86) | Female | 67 | White-European<sup>a,b</sup> |
| FECD6exp+ (12/106) | Male | 69 | White-European<sup>a,b</sup> |
| FECD7exp+ (16/94) | Female | 54 | White-European<sup>a,b</sup> |
| FECD1exp- (13/18) | Female | 64 | European/Black-Caribbean<sup>a</sup><br>Black-African<sup>b</sup> |
| FECD2exp- (18/24) | Female | 73 | White-European<sup>a,b</sup> |
| FECD3exp- (22/28) | Male | 53 | Black-African<sup>a,b</sup> |
| *Spatial transcriptomics* |  |  |  |
| Control 8 (12/16) | Female | 45 | Black-African<sup>a</sup> |
| Control 9 (15/33) | Female | 46 | White-European<sup>a</sup> |
| Control 10 (25/26) | Male | 70 | White-European<sup>a</sup> |
| FECD4exp- (15/15) | Female | 48 | White-European<sup>a</sup> |
| FECD5exp- (16/23) | Female | 57 | Black-African<sup>a</sup> |
| FECD6exp- (15/16) | Female | 58 | Black-African<sup>a</sup> |
| FECD8exp+ (63/88) | Female | 75 | White-European<sup>a</sup> |
| FECD9exp+ (12/76) | Male | 61 | Black-African<sup>a</sup> |
| FECD10exp+ (18/67) | Male | 65 | White-European<sup>a</sup> |

CTG18.1 expansion status is denoted in brackets for all FECD-case derived CEC lines analysed. \*Age at time of corneal tissue excision. <sup>a</sup> self-reported by affected individual. <sup>b</sup> ancestry determined by applying a principal component analysis (PCA) method, FRAPOSA (22). All control samples were derived from donor corneas with very high CEC counts ( $\geq 2,600$ cells/mm<sup>2</sup>) and endothelium free of guttae or any other abnormalities.

**Table 2:**
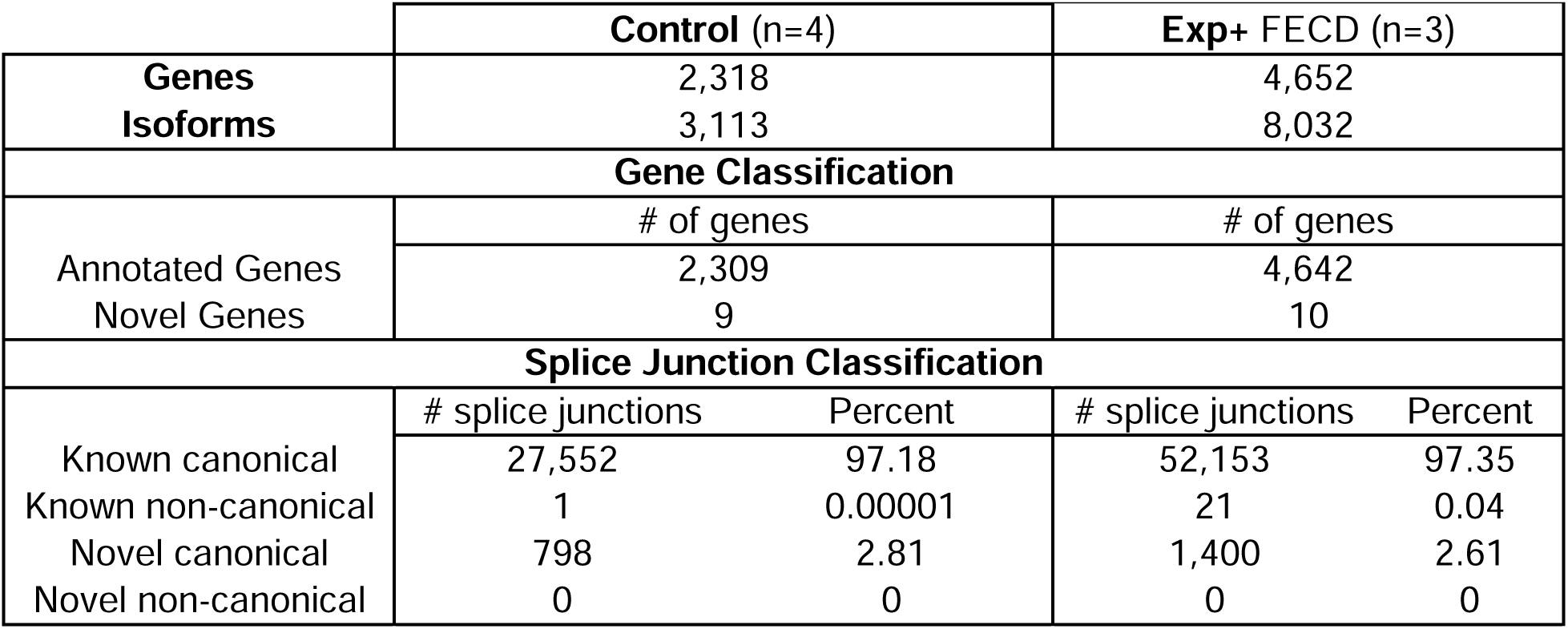
Summary of SQANTI3 analysis of IsoSeq data. In-depth characterisation of isoforms in control and Exp+ long-read RNA-seq data utilising SQANTI3 pipeline following stringent quality control. SQANTI3 subdivides splice junctions into known (present in reference transcriptome) and novel (not in reference transcriptome). Canonical splice junctions refer to the most common type of splice junction (GTAG, GCAG, ATAC)), while non-canonical refers to other rarer and tissue specific splice junctions with different donor and acceptor sites. SQANTI3 defines novel as genes or splice junctions that are absent from the reference transcriptome.

### Alternative splicing analysis of long-read and short-read transcriptomic data demonstrate the complexity of increased splicing events underlying CTG18.1 expansion-mediated disease in CECs

Iso-seq and subsequent SQANTI3 analysis (**Table 2, S3**) revealed a higher proportion of isoforms per gene in Exp+ samples (1.7 unique isoforms per unique gene in Exp+ vs 1.3 in control). Additionally, we see a higher proportion of known non-canonical splice junctions in Exp+ samples. When characterising transcripts based on splice junctions, we see similar proportions of isoform subtypes (**Table S3**). This is consistent with previous work (6,10–12) and validated by our short read data, discussed below, (**Figure S1, S2**), whereby Exp+ samples showed greater isoform diversity compared to control and Exp- samples, but not an overall skew in the types of alternative splicing observed.

To further characterise and quantify alternative splicing events within our short-read dataset, we utilised rMATS, a junction-based method that employs a hierarchical framework, to model unpaired datasets and evaluate and quantify all major alternative splice types (23–25). Comparing Exp+ to control samples (PWC1) we identified 2,118 significant alternative splicing events in 1,437 genes (**Figures S1, S2, Table S4)**. Alternative splicing comparisons including Exp+ (PWC1 and PWC2) yielded the most alternative splice types in every category (**Figure S1, S2**) further reinforcing the spliceopathy aspects of this genetic subtype of disease.

We then compared the rMATS and Iso-seq results to a subset of published datasets exploring differential splicing in FECD samples (**Table S5**) and selected 24 published differential splicing events (from 23 genes) that had previously been identified by bioinformatic or *in vitro* methods to be strongly associated with CTG18.1 expansion-mediated FECD (4,6,7). Alternative splicing analysis of our transcriptomic dataset identified that 75.0% (18/24) of these events were significant in PWC1 (control vs Exp+, **Table S5**). In addition to these 24 selected events, rMATS identified another 10 novel and significant differential splicing events within Exp+ in these genes, highlighting the increasing complexity of splicing dysregulation within Exp+ FECD. Long-read sequencing confirmed the presence of these novel splicing events in Exp+ FECD (**Table S6**). Also, it was interesting to note that *FN1* was identified to have robust alternative splicing in Exp+, and further investigation showed that alternatively spliced exons in Exp+ corresponded with skipping of the Extra Domain A (EDA) and Extra Domain B (EDB) protein domains that play a critical role in signal transduction and fibrillar formation (26) (**Table S7**).

Skipped exon alternative splicing events, where an additional exon is either included or excluded, were the most common in all three pairwise comparisons (**Figure S1**). Prior work regarding alternative splicing in FECD has focused on MBNL1/2 sequestration(6,7,27) as MBNL proteins are key regulators of exon splicing(28,29). Given their complex role, we attempted to determine if the flanking intronic sequences of alternatively spliced exons were enriched for common RNA-binding protein (RBP) binding motifs. We focussed our RBP motif enrichment analysis (rMAPS2(30)) on the 250bp region upstream of alternatively spliced exons and isolated those areas where there was enrichment in PWC1 and PWC2, but not PWC3 (i.e. events enriched within Exp+ CEC lines). By filtering rMAPS results in this manner, a trend whereby skipped exons are significantly (p < 0.05) upregulated by RNA-binding proteins with TT-containing binding motifs and downregulated by RNA-binding proteins with CG-containing binding motifs was observed (**Table S8**). As MBNL proteins favour YCGY binding motifs, this suggests that other RNA-binding proteins with TT-containing motifs could also play a role in driving the spliceopathy underpinning RNA toxicity, a key mechanism of disease in Exp+ FECD.

### Evidence of TCF4 dysregulation appears enriched in CTG18.1-mediated FECD CECs, highlighting specific candidate biomarkers for this genetic subtype of disease

Differential gene expression was assessed with DESeq2(31) in all three groups including sex and ethnicity as covariates. For all comparisons an FDR corrected P-value (padj) of <0.05 was used as a threshold for significance (**Table S9**). A total of 4,288 genes were identified to be differentially expressed between the Exp+ and controls (PWC1; **Figure 2A**). Of these, 2,089 genes had a shrunkLFC value (moderated log_2_fold values of expression accounting for high dispersion and low expression counts) of greater than 1 or less than -1. Overall, PWC2 and PWC3 groups resulted in lowered differential gene expression, with 1,695 and 1,911 significantly differentially expressed genes, respectively **(Figure 2A; Table S9)**.

**Figure 2:**
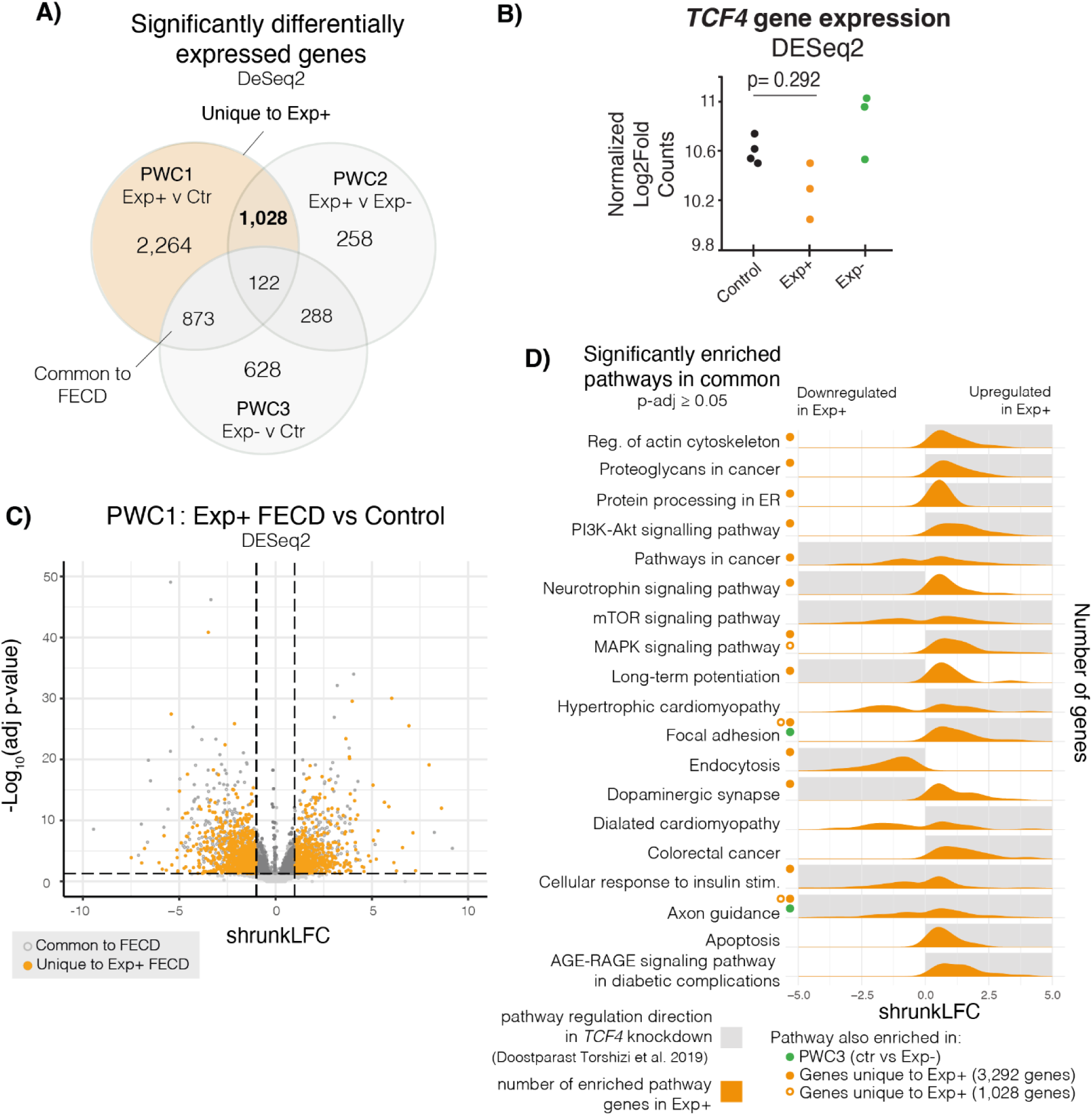
Differential gene expression analysis on Exp+ FECD, Exp- FECD, and control primary corneal endothelial cell (CEC) transcriptomes. **A)** Differential gene expression was assessed via DESeq2. Three pairwise comparisons were conducted: PWC1 (Exp+ vs control), PWC2 (Exp+ vs Exp-), and PWC3 (Exp- vs control). Venn diagrams show overlap of significantly differentially expressed genes (adjusted p-value < 0.05) within these three comparisons. **B)** Total *TCF4* expression showed no significant dysregulation between the different groups. **C)** Volcano plot shows the distribution of gene expression in Exp+ FECD compared to the control group. Horizontal dotted lines denote the boundary between significance (adjusted p-value < 0.05), whereas vertical dotted lines show a shrunkLFC cutoffs for -1 and 1. Dark grey solid markers show significantly expressed genes with a fold change magnitude greater than 1. Solid orange markers delineate uniquely Exp+ differentially expressed genes. **D)** Pathways enriched in Exp+ vs Control (g:profiler). Shaded grey boxes show direction of pathway enrichment in TCF4 knockdown models (Doostparast Torshizi et al. 2019). Green dots show pathways that are significantly enriched in PWC3 as well. Solid orange dots and open orange circle show that pathway enrichment persists in genes unique to Exp+ (1,028 genes and 3,292 genes, respectively)

Pairwise analysis revealed considerable overlap between dysregulated genes identified by all three pairwise comparisons **(Figure 2A),** however, highest convergence was observed by PWC1 (Exp+ versus control) and PWC2 (Exp+ v Exp-), suggesting these events are driven by CTG18.1 expansions. In total, both pairwise comparisons reveal 1,028 dysregulated genes uniquely and significantly dysregulated in Exp+ FECD case-derived samples. Conversely, by comparing the overlap between PWC1 and PWC3 (Exp+ vs control and Exp- vs control) we identified a further 873 genes that are dysregulated in FECD, irrespective of the underlying genetic cause, and thus representing more ubiquitous signatures of the disease. Dysregulated genes in PWC1, including Exp+ unique and common to FECD, did not skew directionally and were evenly up or down-regulated within our analysis (**Figure 2C**). We isolated 3,292 genes that are uniquely dysregulated in Exp+ FECD, 2,264 genes dysregulated solely in PWC1 and a further 1,028 dysregulated genes shared with PWC2; (highlighted in orange **Figure 2A**) and proceeded with downstream analysis (**Table S9**).

We conducted enrichment analysis using g:profiler (32) (querying gene ontology (GO), KEGG pathway, and Reactome pathways). GO term enrichment and KEGG pathway analysis of PWC1 dysregulated genes was consistent with previously published transcriptomes (6,11,12), with several enriched terms and pathways relevant to RNA toxicity (i.e. GO:0000166 etc) (**Table S10**). While total *TCF4* gene expression was not significantly downregulated in Exp+ (**Figure 2B**), intriguingly, we observed strong parallels to pathway enrichment analyses highlighted from a study of *TCF4* neural progenitor knockdown cell model (33) within the 3,292 genes uniquely dysregulated in Exp+, such as protein processing in endoplasmic reticulum, MAPK signalling, actin cytoskeletal regulation, endocytosis and apoptosis (**Figure 2D**). Critically, these pathways are not enriched in PWC3, suggesting that these are unique to Exp+ CECs. Furthermore, several enriched pathways and GO terms involving Wnt signalling were seen in the 3,292 dysregulated genes unique to Exp+ that are not present in PWC3, again suggesting *TCF4* function may be modified within Exp+ FECD CECs (**Table S10**). Furthermore, enrichment of dysregulated genes in pathways involving other repeat expansion-mediated diseases and misfolded protein aggregate diseases suggest mechanistic parallels which reinforce Exp+ CECs as a model system with broad relevance to repeatome biology (**Table S10**).

The consistent presence of PI3K-Akt pathway enrichment within KEGG results in Exp+ dysregulated genes (and its consistent absence in PWC3 pathway enrichment) suggests that Exp+ FECD CECs may interact with growth factors and the extracellular matrix in a unique manner. Hence, components of the PI3K-Akt pathway may serve as informative biomarkers for Exp+ FECD in future clinical trial settings, and thus warrant further investigation. Additional pathways are enriched in this set of genes (and not in PWC3) that involve RNA processing and binding, other repeat-mediated disease related mechanisms, epithelial to mesenchymal transition, and multiple signal transduction cascades with extracellular matrix interactions (**Table S10**).

To determine if other RNA-binding proteins may be implicated in the alternative splicing seen in Exp+ FECD, we cross-referenced our short-read data with a list of ∼1500 human RNA-binding proteins (34), and discovered that there were 141 RBPs uniquely dysregulated in Exp+ (45 downregulated and 106 upregulated), including *MBNL1, CELF1, ESRP2,* and a variety of hnRNP and DDX encoding genes (**Table S11**). Both the hnRNP and DDX family of RNA-binding proteins have previously been implicated in other repeat expansion diseases (32,35–37). It is likely that a combination of these dysregulated RBPs plays a role in driving the alternative splicing and RNA toxicity seen in Exp+ FECD.

### Differential TCF4 exon usage suggests isoform ratio disruption of TCF4 in CECs of CTG18.1-mediated FECD individuals

Given signatures of *TCF4* dysregulation were apparent in the pathway enrichment analysis of Exp+ CECs (**Figure 2D**), we then aimed to determine if any tangible patterns of *TCF4* transcript-specific dysregulation was observed in the Exp+ CECs, despite total *TCF4* levels not being significantly different between Exp+, Exp- and control CECs **(Figure 2B).** Due to transcriptional diversity and complexity of TCF4 transcripts, DEXSeq (38) was used to assess differential exon usage utilising our short-read transcriptomic data.

DEXSeq utilises generalised linear models to detect differential exon usage while controlling for biological variation. This analysis allowed us to query gene expression and splicing at the exon level comparing PWC1 (with and without (PWC1_nocov) sex and ethnicity as covariates), PWC2, PWC3, FECD compared to control (FvC) and Exp+ compared to all expansion negative samples (Exp+ v Ctr+Exp-). In concordance with our rMATs and DESeq2 analysis, higher levels of transcriptome-wide differential exon usage events were observed in PWC1 compared to PWC3 (**Figure S3**), suggesting these patterns of differential exon usage are likely attributed to CTG18.1-related pathology.

Our short-read transcriptomic data reveals that multiple *TCF4* exons (n=5) were significantly dysregulated in Exp+ compared to control and Exp- FECD samples (**Figure 3A, Table S12 and S13**), whereas no significant differences were observed between Exp- and control (PWC3, **Figure 3B, Table S12 and S13**), demonstrating for the first time that differential *TCF4* exon usage is driven by CTG18.1 expansions and not a downstream feature of FECD unrelated to the primary genetic driver. A subset of exons contained within a small group of *TCF4* transcripts (4 out of 93) are upregulated in Exp+, while an increased number of more broadly expressed exons are downregulated (appearing in 56 out of 93 transcripts) (**Figure 3A, Table S12 and S13**). Transcripts containing downregulated exons are primarily included in longer isoforms of the gene which contain 3 AD domains, while the upregulated exons (excluding those surrounding the repeat) appear in shorter *TCF4* isoforms containing only two AD domains. This suggests that Exp+ CECs may be employing certain compensatory measures in response to the transcript-specific dysregulation that is induced by the presence of CTG18.1 repeat expansions. Interestingly, our rMATS analysis independently detects this same phenomenon (**Table S14**), which showed a significant mutually exclusive exon splicing event whereby the last exon in common with most 3 AD domain-containing isoforms (Exon 6 in ENST000000354452.8) was favoured in the control group, while the first exon in common with all isoforms (Exon 7 in ENST000000354452.8) was favoured in the Exp+ group.

**Figure 3:**
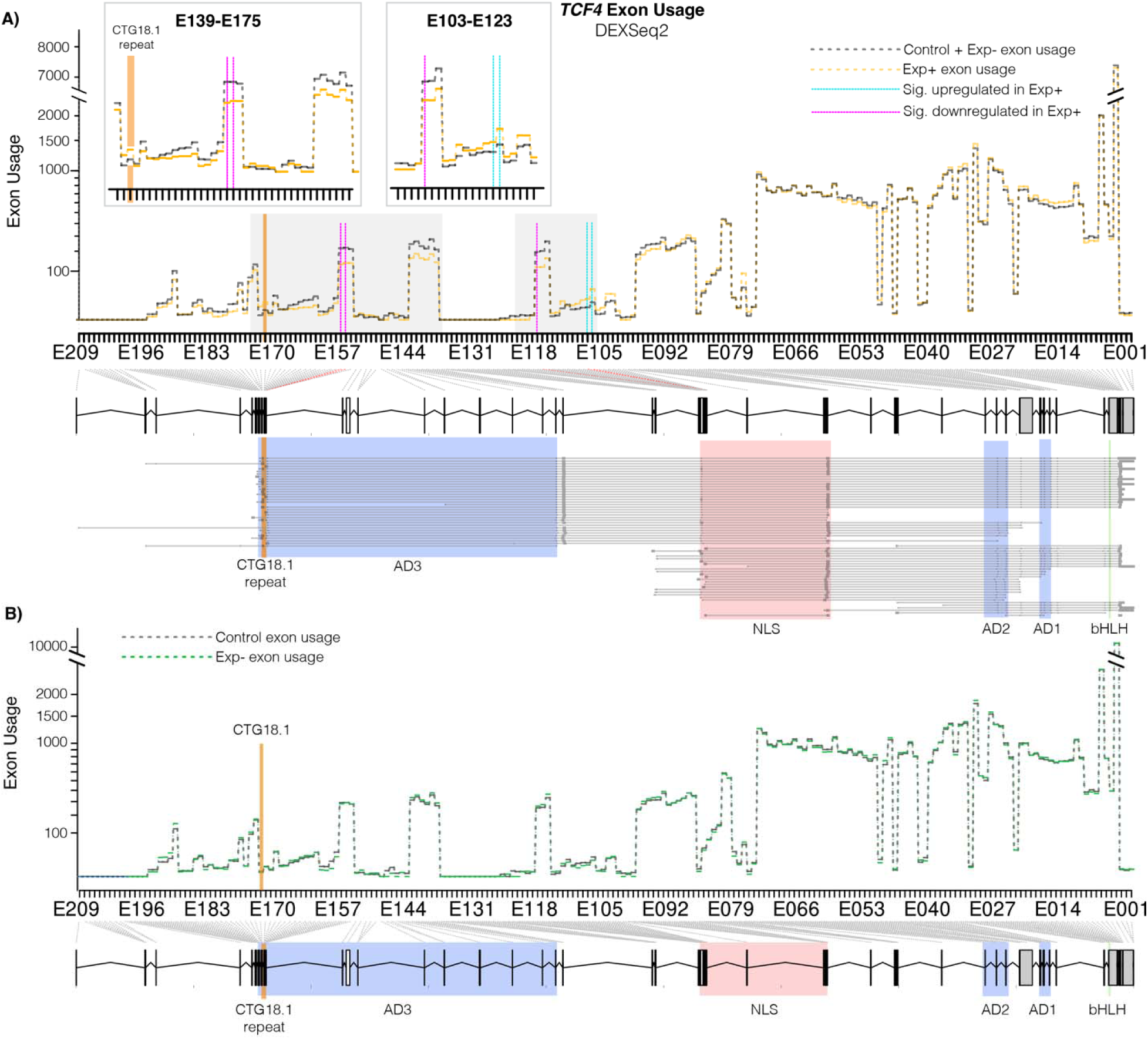
CTG18.1 expansion-mediated FECD results in alternative *TCF4* exon usage within corneal endothelial cells (CECs). **A)** Differences in exon usage between control (grey dotted line) and Exp+ (orange dotted line) CECs. Significantly upregulated and downregulated exons are highlighted with blue and red vertical lines, respectively. The location of CTG18.1 is depicted with an orange line. Inset boxes magnify dysregulated regions. A genomic schematic of *TCF4* with transcripts aligned is presented below the axis. Key protein domains highlighted (green - basic helix loop helix (bHLH) domain, blue - activation domains (ADs), red - main nuclear localization signal (NLS). **B)** Differences in exon usage between control (grey dotted line) and Exp- (green dotted line) CECs. No exons were statistically dysregulated.

We also detect retention of the intron flanking the CTG18.1 repeat, previously described by Sznajder et al. 2018 (39) in Exp+ FECD **(Table S12 and S13).** The intronic region is significantly upregulated in Exp+ CECs (E173 and E174) **(Table S12 and S13)**. While this intron represents the 5’ untranslated region (UTR) of several transcripts of *TCF4*, it is also included within an intronic region in 32 out of 93 currently annotated *TCF4* transcripts.

### RNAScope shows robust detection of altered tissue-specific TCF4 isoform expression in CTG18.1-mediated FECD case-derived corneal endothelial cells

We performed spatial transcriptomics, via RNAScope, to further validate our hypothesis that a subset of *TCF4* isoforms containing the third AD3 domain are being selectively downregulated due to the presence of an expanded CTG18.1 allele (≥50 repeats). For the first time, we visualised *TCF4* transcripts within FECD and control CECs and fibroblasts using two distinct sets of probes, termed A and B (**Figure 4A**). Probe set A comprises a set of probes that allow the visualisation of all *TCF4* transcripts in the cell (probe A covers all exons in the transcript Isoform B+ which contains 3 AD domains, TCF4-201, ENST00000354452.8, NM_001083962.2). Probe set B targets exons in the canonical longer TCF4-201 *TCF4* transcript which do not overlap with the canonical shorter *TCF4* transcript (Isoform A+/J which contains 2 AD domains, AD1 and AD2, TCF4-204, ENST00000457482.7, NM_001243234.2). Hence, these probe sets enable the detection, visualisation, and quantification of the ratio of longer AD3-containing transcripts, previously identified to be downregulated in Exp+ CECs via our differential exon usage analysis, compared to total *TCF4* transcripts (**Figure 3A**).

**Figure 4:**
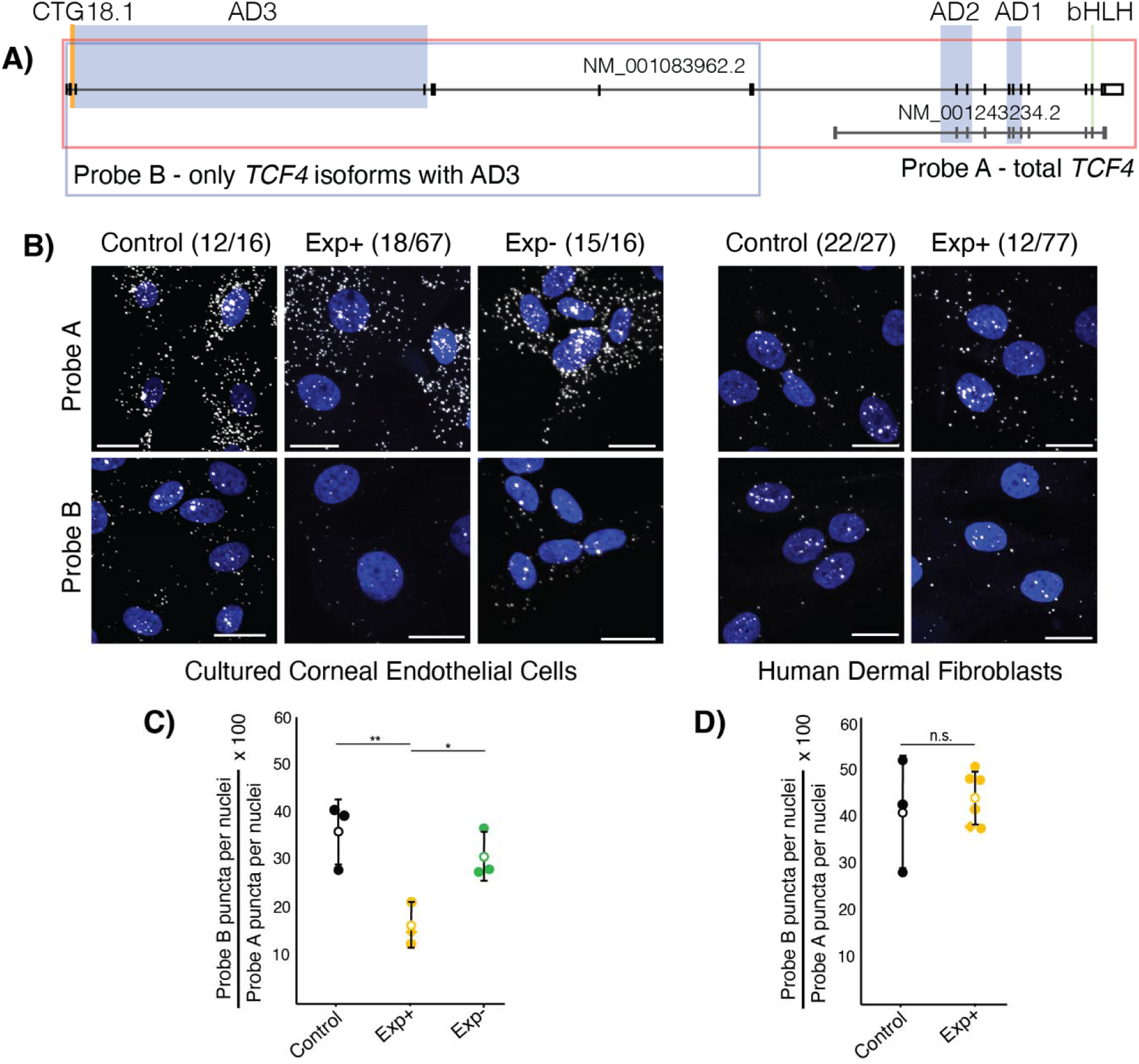
RNAScope probes targeting *TCF4* transcripts demonstrate that longer transcripts containing activation domain 3 (AD3) account for a smaller proportion of total transcripts specifically in Exp+ FECD corneal endothelial cells (CECs). A) Probe targeting schematic for *TCF4*. Probe A targets all exons in NM_001083962.2 and visualises total *TCF4* transcripts, while Probe B targets only the 5’ exons, excluding those that are contained within shorter transcripts (NM_001243234.2). Probe B visualises the longer subset of AD3-containing transcripts within the cell. CTG18.1 is highlighted in orange. Green bar denotes TCF4 bHLH region, blue regions highlight TCF4 activation domains, red box shows the bipartite TCF4 NLS signal and orange shows the genomic region containing the CTG18.1 repeat. Key domains highlighted (green - basic helix loop helix (bHLH) domain, blue - activation domains (ADs), red - main nuclear localization signal (NLS)). **B)** Representative confocal images of Exp+, Exp- and control CECs and Exp+ and control fibroblasts with nuclei (blue, DAPI) and *TCF4* puncta (visualised with Cy3 but represented here with white). Scale bars represent 20 µm. **C)** Plotting the proportion of longer AD3-containing *TCF4* transcripts in Exp+ (orange), control (black), and Exp- (green) shows that Exp+ have significantly lower proportion of AD3-containing transcripts compared to both control (p = 0.0145) and Exp- (p = 0.0213) CECs, while there is no significant difference between control and Exp- (p = 0.362). D) Plotting the proportion of longer AD3- containing *TCF4* transcripts in Exp+ (orange), and control (black) shows that there is no statistically significant (p=0.61) difference in *TCF4* isoform ratios between the two groups. Open circle denotes mean and error bars show standard deviation. Diamond datapoint shows bi-allelic expanded FECD CECs. Significance determined with student T-test.

The *TCF4* transcripts were visualised in control, Exp+ and Exp- FECD case-derived CECs (n=3 for each group) (**Figure 4B**, **Table 1, S1 and S15)**. Each independent primary CEC culture was also probed with a negative control (targeting bacterial genes) and a (CAG)_7_-Cy3 probe, using a standard FISH protocol, to validate the presence or absence of CUG-specific RNA foci derived from expanded copies of CTG18.1 (**Table S15**). RNA foci were consistently detected in all CTG18.1 Exp+ CECs, and were absent from control and Exp- CECs. Using CellProfiler (version 4), puncta representing *TCF4* transcripts, detected with both probe set A and B, were quantified and puncta/cell values were established for each image and averaged across images with at least 100 nuclei imaged per condition per line. Finally, a ratio was calculated establishing the proportion of longer AD3 containing transcripts compared to total *TCF4* transcripts for each biological replicate. This showed that, on average, Exp+ CECs had a significantly smaller proportion of the longer AD3-containing transcripts (16.33% ± 4.53) compared to unaffected control CECs (36% ± 6.9, p=0.0145) and non-expanded FECD CECs (30.9% ± 5.17, p=0.0213) (**Figure 4C**). To explore potential cell-type specificity of the observed skewed *TCF4* transcript ratios we also analysed a series of dermal fibroblast lines generated from unrelated Exp+ patients with FECD (n=6) (7) and control adult dermal fibroblast lines (n=3) (Table S16). No significant difference (p=0.61) was observed between the Exp+ (44.2% ± 5.68) and control fibroblast (41.17% ± 12.1) lines, suggesting the dysregulation of *TCF4* is cell-type specific. This result suggests that the presence of the expanded repeat alone consistently skews *TCF4* transcript ratios to favour shorter isoforms, specifically in CECs. As this subgroup of isoforms typically contains only two out of three activation domains, a higher prevalence of shorter TCF4 isoforms may affect potential dimerization and TCF4 transcriptional function in Exp+ CECs.

### Rare TCF4 variants identified within a genetically refined CTG18.1 expansion-negative FECD cohort suggest isoform-specific TCF4 dysregulation may be a risk factors for FECD in the absence of CTG18.1 expansions

We next wanted to explore if rare and potentially deleterious *TCF4* variants could be contributing to disease in FECD cases without a CTG18.1 expansion, given that Exp+ CECs display alternative *TCF4* exon usage and skewed ratios of *TCF4* isoforms (**Figure 3 and 4**), with detectable downstream features of TCF4-driven dysregulation (**Figure 2D**). To test this hypothesis, we interrogated exome data generated from 134 Exp-FECD cases, acquired from our ongoing inherited corneal disease research program, representing approximately 22% of total FECD cases recruited to date. Overall, 8 rare and potentially deleterious (MAF <0.005; CADD score >15) *TCF4* variants were identified in 7/134 probands (**Table 3**). Sanger sequencing confirmed the presence of all 8 variants and exome data from these cases were also interrogated using the same thresholds (MAF <0.005, CADD >15) to determine if any other rare variants in genes previously associated with FECD could also be contributing to disease (4) (**Table S17**). The large number of rare and potentially deleterious *TCF4* variants detected in this cohort seemed intriguing given the high levels of evolutionary constraint on this essential and ubiquitously expressed transcription factor encoding gene (pLI=1, missense Z=4.1 gnomAD). We therefore decided to apply CoCoRV, a rare variant analysis framework to determine if the Exp-FECD cohort was indeed enriched for rare and potentially deleterious *TCF4* variants compared to the gnomAD dataset. CoCoRV utilises the publicly available genotype summary counts to prioritise disease-predisposition genes in case cohorts (40). Under a dominant model, applying the ethnically-stratified (**Table S18**) analysis pipeline, and filtering for rare (MAF <0.005) and potentially deleterious (CADD score threshold of >15) variants, an inflation factor estimate (λemp) 1.21 was achieved. At the exome wide level *TCF4* was ranked as 34/17,025 genes with a p-value of 0.001 suggesting the Exp-FECD cohort could be enriched for rare and potentially deleterious *TCF4* variants. However, after applying CoCoRV’s stringent false discovery rate correction methodology *TCF4* no longer remained significant (corrected p-value = 0.249).

**Table 3:** Summary of rare and potentially deleterious *TCF4* variants identified in a FECD CTG18.1 Exp- cohort (n=134)

| Variant ID | FECD Exp-Cases (n=134) | Variant (hg38) | HGVSc, HGVSp | CADD, MAF gnomAD | Included within CoCoRV analysis |
| --- | --- | --- | --- | --- | --- |
| 1a | Proband A | chr18:55588479-C-A | ENST00000566286.1: c.57G>T, p.(Arg19Ser) | 18.3, 0 | Yes |
| 1b | Proband A | chr18:55588478-T-A | ENST00000566286.1: c.58A>T, p.(Lys20Ter) | 16.45, 0 | Yes |
| 2 | Proband B | chr18:55588470-C-T | ENST00000566286.1: c.66G>A, p.(Glu22=) | 18.39, 0.002122 | Yes |
| 3 | Proband C | chr18:55464063-T-C | ENST00000354452.3: c.207+13A>G | 20.6, 0.0002234 | Yes |
| 4 | Proband D | chr18:55403689-A-G | ENST00000544241.2: c.26T>C, p.(Ile9Thr) | 17.29, 0 | Yes |
| 5 | Proband E | chr18:55321599-C-T | ENST00000354452.3: c.549+28760G>A | 16.74, 0 | No* |
| 6 | Proband F | chr18:55261560-A-C | ENST00000354452.3: c.923-27T>G | 22.3, 0.0007644 | Yes |
| 7 | Proband G | chr18:55261512-G-A | ENST00000354452.3: c.944C>T, p.(Ala315Val) | 24.6, 0.0005734 | Yes |

Rare (MAF < 0.005 in gnomAD genomes) and potentially deleterious (CADD score threshold of >15) *TCF4* variants identified within a FECD CTG18.1 Exp- cohort (n=134). Abbreviations are as follows: gnomAD, The Genome Aggregation Database; MAF, minor allele frequency, HGVSc, coding DNA sequence based on the Human Genome Variation Society; HGVSp, protein sequence based on the Human Genome Variation Society. \*Variant filtered out at Variant Quality Score Recalibration step.

Interestingly rare variants identified in Proband A (Variant 1a/1b), B (Variant 2) and D (Variant 4) and E (Variant 5) are all predicted to only affect a small fraction of total *TCF4* transcripts (**Table 3, S19 and Figure 5**). For Proband A, two consecutive *in cis* heterozygous missense and nonsense variants were identified. These occur within the coding region of only one (out of 93) *TCF4* transcripts (ENST00000566286.5; c.[57G>T; 58A>T]; p.[(Arg19Ser; p.Lys20*)]) and in the 5’ UTR of a further 5/93 transcripts. The variant identified in Proband B is only encompassed by the same 6 *TCF4* transcripts (ENST00000566286.5; c.66G>A, p.(Glu22=)). Despite being synonymous, Splice AI predicts the variant introduces loss of the splice donor site (SpliceAI Δ score 0.78). Furthermore, SpliceRover also predicts that this could result in the activation of a cryptic splice donor site downstream (from 0.098 to 0.233) of the wildtype donor site, which would introduce a short frameshift insertion followed by a premature termination codon (PTC) c.66_67insGTGCTCGATGAATTTTC, p.(Arg23Valfs*12). The variant identified in Proband D (Variant 4) is within the protein coding region of three (out of 93) *TCF4* transcripts and in the 5’ UTR of one additional transcript. Variant 5 is only encompassed by the 5’ UTR of a single *TCF4* transcript. Collectively, all of these identified variants have the potential to exert a functional impact on TCF4, but importantly only on a small subset of overall transcripts.

**Figure 5:**
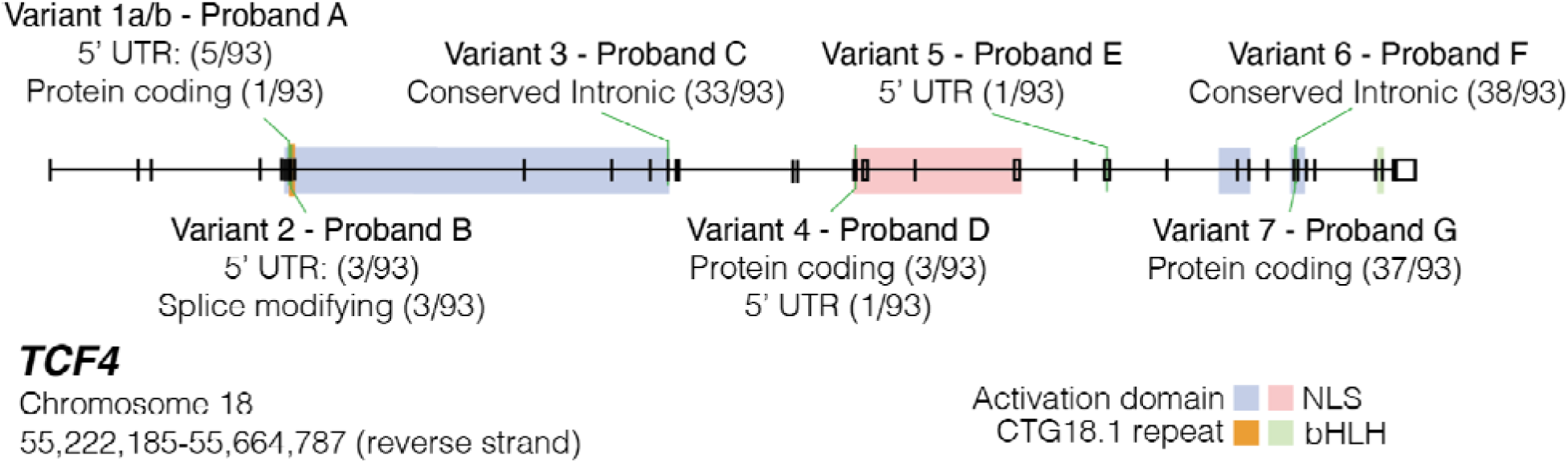
Schematic map of rare and potentially deleterious variants identified in individuals with molecularly unsolved FECD within *TCF4*, without CTG18.1 expansions. All 8 rare (minor allele frequency (MAF)<0.005) and potentially pathogenic (CADD>15) variants in 7 Exp- Probands are mapped along *TCF4* genomic region and transcriptomic locations contextualised. Proband A variants are found in *cis* within the proband. Conserved intronic transcripts refer to variants located in intronic regions identified as conserved via Genomic Evolutionary Rate Profiling (GERP) conservation scores on Ensembl. Splice modifying refers to transcripts where the variant identified would induce alternative splicing. Transcript numbers where the variant occurred within introns were not included. UTR: untranslated region. NLS: nuclear localization signal. bHLH: basic helix loop helix

## Discussion

FECD is now recognized to be the most common repeat expansion-mediated disease in humans, with up to 80% of European FECD cases harbouring CTG18.1 expansions(4). The non-coding nature of the repeat and the transcriptomic complexity of *TCF4* have made it extremely experimentally challenging to decipher the molecular mechanisms underlying this disease. Thus, many aspects of its pathophysiology remain elusive, despite the strong mechanistic parallels with other repeat-mediated diseases, and the urgent clinical need for new treatments for this common and visually disabling disease. To tackle this challenge, we employed a large, biologically independent set of primary FECD case-derived CEC lines and numerous complementary transcriptomic methodologies to gain novel mechanistic insight into both RNA toxicity driven and *TCF4* dysregulation-related molecular features of this repeat-mediated disease. By including both Exp+ and Exp- FECD data, we have been able to dissect out more generic downstream biomarkers that occur irrespective of CTG18.1 expansion status. Our extensive long- and short-read RNA-Seq data and spatial transcriptomic analysis of FECD case-derived CEC lines demonstrates that, in addition to being a spliceopathy, expansion-positive FECD is concurrently associated with isoform-specific patterns of *TCF4* dysregulation, hence providing novel mechanistic insights into the pathophysiology of this triplet-repeat mediated disease. Furthermore, we have identified a subset of Exp- FECD cases with rare and potentially deleterious *TCF4* variants, supporting the hypothesis that dysregulation of TCF4 itself irrespective of CTG18.1 expansions may also be a risk factor for FECD.

Our robust transcriptomic analysis, presented within a biologically relevant cellular and genomic context, highlights an array of novel molecular signatures driven by CTG18.1 expansions, as well as validating a range of previously described Exp+-associated transcriptomic events. We anticipate that many of the novel events identified here could serve as informative biomarkers to aid the development of CTG18.1 targeted therapeutic approaches. To date, CUG-containing RNA foci and MBNL1/2 missplicing have been adopted as the standard for therapeutic outcome measures (7,27,41,42). Data presented here support and expand the hypothesis that accumulation of repeat expansion-derived RNA structures induce toxic gain-of-function effects that disrupt cellular homeostasis and induce aberrant RNA metabolism(7,27). For instance, we have discovered a novel array of RBP encoding transcripts, including *MBNL3*, *CELF1* and a range of *hnRNP* and *DDX* family members, uniquely dysregulated in Exp+ CECs that warrant future follow up to elucidate their respective contributions to aberrant RNA metabolism related mechanisms. However, such biomarkers may only relate to the spliceopathy aspects of the disease, and our data go on to also illustrate that translational efforts would also benefit from using more diverse outcome measures. In addition to RNA metabolism biomarkers, global Exp+-mediated differential gene expression profiles, alongside *TCF4*-specific dysregulation itself, should also be included as relevant outcome measures for Exp+ therapies being developed, given we hypothesise these processes are also pivotal to the pathology. Novel cell surface biomarkers reported here that are specifically upregulated in Exp+ (e.g. *ITGA5* and *ITGB1*) could have particular utility in a clinical trial setting due to their accessible locations. Other membrane encoding proteins interfacing with the extracellular matrix or signal transduction could also be a rich source of potential external markers, such as cell surface receptors involved in the P13K-Akt pathway (e.g. *EGFR, MET,* and *LTK*) that we show are specifically upregulated in Exp+ CECs.

Genetic stratification of patient-derived CECs in this study has also highlighted that Exp+ FECD is transcriptionally distinct from Exp- disease. The data show that 76% of all differentially expressed genes, 81% of skipped exon events, and 67% of all enriched pathways detected in Exp+ are unique to the repeat expansion disease. For example, in accordance with previous studies (12,43), we observe that pronounced upregulation of fibronectin (*FN1*) is a robust transcriptomic feature of FECD, irrespective of genetic underlying cause. However, our comprehensive transcriptomic analysis has enabled us to determine that the EDA/B+ forms of fibronectin (26) are significantly favoured in Exp+ FECD. This illustrates that *FN1* expression and splicing can act as both a general FECD molecular biomarker and an Exp+ specific biomarker. Similarly, *AQP1* previously documented to be downregulated in FECD (44), has here been demonstrated to only display marked downregulation in Exp+ CECs, illustrating the importance of stratifying cases based on CTG18.1 status. Our long-read data suggests that the splicing diversity seen in Exp+ FECD is not predominantly a result of abnormal missplicing, given the proportion of annotated full splice match transcripts remains the same between control and Exp+. This further exemplifies the unique transcriptome of Exp+ FECD endothelial cells. However, the increased proportion of non-canonical splice junctions seen in Exp+ also suggests that CTG18.1-specific isoforms may be present within the endothelium and warrants future investigation. However, transcriptome wide differences in coverage were observed between the Exp+ FECD versus the control group within the long-read data, thus it will be important for future long-read approaches to further validate these findings. These examples, alongside the emerging evidence that the Exp+ FECD displays distinctive pathological features and epidemiological characteristics(5,7), suggests that Exp+ FECD should be categorised as a distinct clinical entity. We suggest integrating CTG18.1 genotyping into routine FECD clinical care pathways is warranted, especially given the numerous international efforts underway to develop gene-directed therapies for CTG18.1-mediated FECD (27,41,42,45).

Our complementary RNA-seq approaches, in combination with the spatial resolution of *TCF4* transcripts subgroups, has highlighted a unique and distinctive pattern of TCF4-specific dysregulation that occurs within Exp+ FECD. Importantly, this signature of dysregulation was not detected within a patient and control dermal fibroblast cell series, implying it may be unique to affected cell types. Additionally, our pathway analysis has isolated features of dysregulation that we hypothesise represent downstream consequences of this isoform shift and an important pathogenic component of the disease. Notably, we have observed commonality between signalling transduction pathways dysregulated in our Exp+ CECs (e.g. P13K-Akt and MAPK) and those dysregulated in neural progenitor cell *TCF4* knockdown transcriptomes(33). We postulate that this convergence is being driven by TCF4 functional dysregulation, as the shift in isoform distributions towards shorter (1-2 AD domains) versus longer (2-3 AD domains) isoforms would result in lowered TCF4 activity within the corneal endothelial cell. TCF4 isoforms with 3 AD domains have been shown to have an order of magnitude more transcription activation function than the shorter isoforms containing 1 or 2 activation domains (16). Furthermore, given that domains in the AD3-containing isoforms can also dictate intracellular location and dimerization partners of TCF4, we also hypothesise that additional TCF4-driven functional implications of this shift occur within affected CECs that requires future experimental investigation. Interestingly, a recent single-cell (sc)RNA-seq study has identified that varying levels of *TCF4* expression exist within discrete populations of healthy CECs (46). It was not possible as part of this study to genotype CTG18.1 whilst concurrently analysing transcriptomic profiles of the CEC populations, thus future development and application of scRNA-seq in combination with single cell genotyping of CTG18.1 within FECD-case derived CEC samples is anticipated to shed light on if and how genomic instability of this repeat element within the corneal endothelium may drive and/or exacerbate the pathophysiology and explain the unique vulnerability of this cell layer to CTG18.1 expansions. (47,48)

Additionally, we identify rare and predicted deleterious *TCF4* variants in 7/134 Exp- FECD cases. This was notable to us given the high levels of evolutionary constraint acting on *TCF4* and suggests such rare variants may be acting as risk factors for FECD, independently of CTG18.1 repeat expansion. This could also in-part explain the established missing heritability of the disease(4) and does further reinforcing the hypothesis that dysregulation of *TCF4* itself is pivotal to the pathophysiology of FECD. Despite a rare variant gene-burden style analysis of these data not reaching significance, we observed an emerging trend whereby a subset of qualifying variants identified notably affected regions of the gene not included by the vast majority of *TCF4* isoforms. Hence, they can only be predicted to exert a functional impact on a small subset of total TCF4 transcripts. This observation is in keeping with the relatively mild and tissue specific nature of FECD, which is in stark contrast to the severe neurodevelopmental disorder Pitt-Hopkins associated with total *TCF4* haploinsufficiency(49). Hence, we propose that *TCF4* variants which subtly affect TCF4 functionality may induce CEC-specific disease. Further experimental investigation of these variants and the potential tissue specific role they may exert in the corneal endothelium warrants further future investigation.

Parallels can be drawn to other repeat expansion-mediated disorders, such as fragile X syndrome (*FRM1*)(50), spinocerebellar ataxia type 7 (*ATXN7,* MIM #164500)(51) and *C9orf72*-mediated ALS(52), whereby rare loss-of-function mutations have been proposed to induce comparable phenotypic outcomes to non-coding repeat expansions. However, in these instances loss of total protein isoform function, in combination with toxic gain-of-function mechanisms, is thought to underlie the respective pathogenic mechanisms, in contrast to this scenario whereby we hypothesise only a fraction of total *TCF4* isoforms are susceptible to the effect of either CTG18.1 expansions or *TCF4* variants within the corneal endothelium. In a limited number of DM1 cases, FECD has been recognised to be part of the phenotypic spectrum, suggesting that CEC dysfunction can also arise from CTG repeat expansions in *DMPK* (53,54). Data presented in this study suggests that *TCF4* variants, independent of the repeat, may also represent a potential path to FECD. We suggest future FECD genetic screening approaches should include sequencing of coding and regulatory regions of *TCF4,* to further elucidate the potential role that rare *TCF4* variants may play in the absence of CTG18.1 expansions.

In conclusion, this study supports the hypothesis that at least two distinct pathogenic mechanisms, RNA toxicity and *TCF4* isoform-specific dysregulation, underpin the pathophysiology of the transcriptionally distinct CTG18.1-mediated FECD. In our established primary corneal endothelial system, we see evidence of *TCF4* isoform dysregulation at the *TCF4* transcript level and can detect the potential downstream effects of the isoform-specific pattern of TCF4 dysregulation. Our data also suggests that, in rare instances, irrespective of CTG18.1 expansions, FECD cases can harbour *TCF4* variants that may independently induce isoform-specific TCF4 dysregulation and act as risk factors for disease. This supports the hypothesis that *TCF4* dysregulation is a key mechanism underlying this common, age-related and visually disabling disease. We anticipate that concepts and biomarkers identified by this study will inform and guide the development of future translational interventions targeting this triplet-repeat mediated disease, and these in turn may act as exemplars for much rarer and currently untreatable and severe repeat expansion disorders that typically affect less accessible neurological/neuromuscular systems.

## Materials and Methods

### Subject Recruitment, phenotyping and bio-sample processing

The study adhered to the tenets of the Declaration of Helsinki and was approved by the Research Ethics Committees of University College London (UCL) (22/EE/0090), Moorfields Eye Hospital (MEH) London (13/LO/1084), or the General University Hospital (GUH) Prague (2/19 GACR). All participants provided written informed consent and had either clinical signs of FECD or had a previous corneal transplant for FECD. For participants who had planned endothelial keratoplasty, we collected the diseased corneal endothelium and Descemet membrane (DM) removed as part of the surgical procedure (MEH). All individuals recruited with FECD undergoing DMEK surgery had confluent guttae affecting the central cornea with visual symptoms and were of similar age. Importantly, all individuals at the time of bio-sample recruitment had similarly advanced disease (Krachmer Grade 5).

We applied strict criteria in the selection for control tissues. We obtained control tissue as donor (age >45 years) corneoscleral discs classed as unsuitable for clinical transplantation due to non-CEC related reasons (Miracles in Sight Eye Bank, Texas, USA). We selected the age of the donor tissues to ensure there was no statistically significant differences in the ages between control and case at the time of bio-sample recruitment. All control corneas were examined and were shown to have a high CEC counts (≥2,600 cells/mm^2^) with no guttae or other endothelial abnormality. To standardise handling, the control DM and corneal endothelium were excised by a consultant-grade surgical clinician in accordance with standardised clinical practice. All excised case and control tissues were stored in Lebovitz L-15 media (Life Technologies) before transferring to the laboratory for processing within 24 hours of excision from either donor corneas or patients undergoing elective posterior corneal transplantation surgery.

Participant genomic DNA for CTG18.1 genotyping was extracted from whole blood (QIAgen Gentra Puregene Blood Kit), whereas control DNA was extracted from the residual corneoscleral disc (QIAgen Blood and Tissue kits).

### CTG18.1 genotyping

We determined the CTG18.1 repeat length using a short tandem repeat (STR) genotyping assay as previously described in Zarouchlioti et al. 2018(7), originally adapted from Wieben et al. 2012(55). Briefly, STR-PCR was performed using a 6’-FAM-conjugated primer (5’ CAGATGAGTTTGGTGTAAGAT 3’) upstream of the repeat and an unlabeled primer (5’ ACAAGCAGAAAGGGGGCTGCAA 3’) downstream. Post-PCR, products were combined with GeneMarker ROX500 ladder and separated on an ABI 3730 Electrophoresis capillary DNA analyzer (Applied Biosystems). Sizing and data analysis was performed on GeneMarker software (SoftGenetics). To distinguish between samples with presumed bi-allelic non-expanded alleles of equal length and samples with a larger repeat length beyond the detection limit of the STR-PCR, we performed a further triplet-primed PCR assay to confirm the presence or absence of a CTG18.1 allele above the detection range of the STR assay (approximately 125 repeats), following previously published methods(56).

### Primary corneal endothelial cell (CEC) culture

We isolated and cultured FECD and control-derived CECs as described by Zarouchlioti et al. 2019(7), adapted from Peh et al. 2015(57). Briefly, to dissolve the DM and detach the CECs, the tissue sample was incubated for 2 to 4 hours in 0.2% collagenase type I. After incubation, the CECs were resuspended in stabilisation media (containing Human Endothelial-SFM (Life technologies) supplemented with 5% foetal bovine serum (FBS), 1% antibiotic/antimycotic, and 0.1% selective ROCK inhibitor Y-27632 (AdooQ BioSciences)), and then seeded into 6-well plates pre-coated with FNC coating mixture (Stratech). After 24 hours, we replaced the culture media with expansion media (Ham’s F-12 Nutrient Mix with GlutaMAX Supplement (Life Technologies)/Medium 199 GlutaMAX Supplement (Life Technologies), 20 μg/mL ascorbic acid, 1% insulin-transferrin-selenium (Life Technologies), 5% FBS, 1% antibiotic/antimycotic, 10 ng/mL bFGF (Life Technologies) and 0.1% selective ROCK inhibitor Y-27632 (AdooQ BioSciences)). All cells were cultured at 37°C and 5% CO_2_ with twice-weekly media changes.

### Fibroblast cell culture

Fibroblast cell lines from dermal skin biopsies were cultured DMEM/F-12, GlutaMAX (Life Technologies) supplemented with 10% FBS and 1% penicillin/streptomycin. Throughout culture, cells were kept in an incubator at 37°C, 5% CO_2_ and medium was refreshed every 48 hr until the cells showed appropriate confluence for experimentation or passage.

### RNA extraction

All primary CEC monolayer cultures selected for downstream analysis displayed a consistent morphology characteristic of CECs *in vivo.* Total RNA was extracted from the primary CEC lines using a NucleoSpin RNA XS kit (Macherey-Nagel), in accordance with the manufacturer’s protocol. The RNA was eluted and stored at -80°C prior to sequencing. To assess RNA integrity, RNA integrity numbers (RIN) were determined for all RNA samples using a Bioanalyzer 2100 (Agilent). All samples analysed as part of this study had a RIN value ≥9.6.

### Short-read RNA-seq

Stranded RNA-seq libraries were prepared by BGI Genomics using the TruSeq RNA Library Prep Kit (Illumina) following the manufacturer’s protocol. We sequenced these with a HiSeq 4000 platform (Illumina) with short-read strand-specific library prep with poly A enrichment (Illumina). Paired-end reads were filtered to remove low-quality reads (Q ≤5) or those with >10% of N base calls. Adaptor sequences were also removed during data quality analysis. We checked the quality of the resulting fastq files for the short-read data using FastQC Quantification(58). Reads were aligned to the human genome (hg38) patch 13 using STAR (59) (v2.7.10a) and Salmon (V1.4.0) (60). After alignment, hierarchical clustering was performed using PoiClaClu(61). We also performed principal component analysis (PCA) using PCAtools(62) (v2.2.0) to determine global sample clustering. Sex and ethnicity have been factored in as covariates in all applicable downstream pipelines.

We analysed differential gene expression using DESeq2(31) (version 1.8.2), with independent hypothesis weighting (IHW) testing of the results to generate FDR values(63,64), and log_2_fold change was calculated using the apeglm (v 1.2) package(65). We then determined significance as FDR corrected-p value (padj) <0.05. In addition, we analysed different exon usage amongst the subgroups with DEXSeq(38) (v 1.44.0). Significant results were filtered using an FDR-corrected P-value of <0.05. Pathway analysis was performed by g:Profiler on significantly dysregulated genes using g:SCS statistics for thresholding (66).

We assessed differential splicing at a junction-based level with rMATS-turbo 4.1.2(23) with the Ensembl hg38 release 105 GTF file as reference. Default rMATS settings were used. For this study, we focused on splicing events that are fully/partly annotated by Ensembl. Significant alternative splicing events were identified as an absolute delta percent spliced in (psi) magnitude ≥ 0.1 and FDR ≤ 0.05. rMATs output was subsequently analysed using rMAPS2(30) and maser 1.12.1(67). All script utilised to generate data can be found at: https://github.com/InheritedCornealDisease/TCF4-transcriptomic-paper-2024

### Long-read RNA-seq

PacBio Sequel long-read transcriptomic sequencing (Iso-Seq)(68) was applied to generate full-length cDNA sequences from control and FECD Exp+ primary CEC-derived RNA samples. Raw reads were processed and de-multiplexed with Iso-Seq (v3) and merged for transcript collapse using Cupcake(68) (parameters: “-c 0.85 -i 0.95 --dun-merge-5-shorter”). High-quality, full-length transcripts from the merged dataset were then aligned to the human reference genome (hg38) using Minimap2(69) (v2.17, parameters: “-ax splice -uf -- secondary=no -C5 -O6,24 -B4”) and annotated using SQANTI3(70) (v7.4). SQANTI filtering of artefacts (intra-priming and reverse transcriptase template switching) was then applied. Github repository for the analysis described here can be found at: https://github.com/SziKayLeung/UCL_FECD

### RNAScope detection of TCF4 transcripts

We seeded primary FECD and control CECs and fibroblasts on chamber slides (Lab-tek) and confluent cell cultures were stained using standard RNAScope protocols with minor adjustments (ACD Bio). We used the Hs-TCF4-C2 (cat. 557411) probe set to detect all *TCF4* isoforms, as well as a custom set of *TCF4* probes to detect all 5’ exons in the *TCF4* transcript NM_001083962.2 excluding all 3’ exons contained in the shorter transcript NM_001243234.2. Akoya TSA Cy3 fluorophores were used to visualise probe staining. Negative control probes (cat. 320871) were provided by ACD-Bio to verify specificity. All cell lines analysed by RNAScope were also tested for RNA foci in parallel via fluorescence *in situ* hybridization (FISH) using a CUG-specific probe (Cy3-(CAG)_7_)(7). A minimum of 100 cells and four images were acquired for each cell line and each probe.

### Exome Sequencing and Rare Variant Analysis

Exome sequencing libraries were generated from 134 whole blood derived gDNA samples from individuals with Exp- FECD, using a SureSelect Human All Exome V6 capture kit (Agilent, USA) or SeqCap EZ MedExome Enrichment Kit (Roche) and sequenced on a HiSeq 4000 or HiSeq 2500 platform (Illumina). All raw FECD case sequencing data was aligned and annotated following our previously published methods(71). In brief, sequencing reads were aligned using Novoalign (version 3.02.08) and variants and indels were called according to Genome Analysis Toolkit (GATK) (version 4.4.0.0) best practices (joint variant calling followed by variant quality score recalibration) were normalised and quality-controlled, as recommended by CoCoRV(40). CoCoRV, a rare variant analysis framework, was applied to prioritise disease-predisposition genes in the Exp-FECD cohort, utilising publicly available genotype summary counts from gnomAD (genomes version 2.1.1) as a control dataset. For consistency, both case and control variants were annotated using the Variant Effect Predictor (VEP) (release 110) (72). The CoCoRV pipeline was then run, applying a dominant model and ethnically stratified in accordance with ethnicity predictions generated from the case cohort using Fraposa(22). Fraposa is based on principal component analysis, using 1000 Genomes project as the reference panel (with known ancestral background, categorised into 5 distinctive superpopulations: Africans, admixed Americans, East Asians, Europeans and South Asians). Common variants between the control and exome cohort were identified and filtered (based on minor allele frequency, linkage equilibrium and missing genotype call rate) by plink and the principal components were computed with the default OADP approach (Online Augmentation, Decomposition, and Procrustes Transformation). Variants of interest were defined as having a CADD score (CADD) >15 (73), and a minor allele frequency (MAF) <0.005 in the Genome Aggregation Database (version 2.1.1) (74). FECD cases identified to harbour rare *TCF4* variants were additionally interrogated for rare and potentially disease-associated variants (MAF<0.005 in gnomAD and Kaviar Genomic Variant Database; CADD>15) in previously reported FECD-associated genes, including *COL8A2*, *SLC4A11*, *AGBL1* and *ZEB1*(4). Splice site predictions were subsequently made using SpliceAI(75) and SpliceRover(76) for all qualifying variants in *TCF4* and any other FECD-associated genes.

## Supporting information

Large Supplemental Tables

## Author Contributions

NB and AED conceptualised the study. NB, NJHT, ANS, CZ, JJ, LD, PS, MP, KM, PL, SJT contributed to the generation of the data. NB, NC, NJHT, AS, SKL, TL, IM, ARJ, MEC, AJH, NP and AED all contributed to the analysis and interpretation of the data. AED and NB wrote the manuscript. All authors read and approved the final version of the document.

## Acknowledgments

We would like to thank all affected individuals for participating in this research. We would also like to acknowledge Miracles for Sight Eye bank for providing control corneal tissue. We would also like to thank Dr. Christopher Smith from Queen Mary University for providing a control fibroblast line. This work was funded by a UKRI Future Leader Fellowship MR/S031820/1 (AED, CZ, AS), Moorfields Eye Charity GR000060, GR001395, GR001337 (AED, NB, NHT, AS, NP) and Sight Research UK SAC 036 (AED, AJS), the Rosetrees Trust (AED, AJS), and the The National Institute for Health Research Biomedical Research Centre at Moorfields Eye Hospital National Health Service Foundation Trust and UCL Institute of Ophthalmology (AED, NC., SJT, KM, AJH, MEC, NP). JJ, LD., PS, and PL were supported by GACR 20-19278S, UNCE/MED/007 and SVV 260631. NP was supported by an NIHR AI Award (AI AWARD02488). This project utilised equipment funded by the UK Medical Research Council (MRC) Clinical Research Infrastructure Initiative (award number MR/M008924/1).

## Supplemental Material Figure and Table legends

**Figure S1: Pairwise comparison of rMATS alternative splicing analysis demonstrates increased levels of alternative splicing in Exp+ FECD compared to both Exp- FECD and control primary corneal endothelial cells.** Short-read RNA-seq data generated from primary corneal endothelial cells were analysed by rMATS. **A)** Table of rMATS events for each pairwise comparison (PWC). Significance denoted by FDR ≤ 0.05 and deltapsi magnitude larger than 0.1. Values in brackets show percentage of total splice events **B)** Venn diagram of skipped exon event coordinates between all PWCs. The largest overlap is observed between PWC1 and PWC2 highlighting an enrichment of skipped exon events in Exp+ FECD compared to Exp- FECD and controls.

**Figure S2: Alternative splicing analyses of short-read CEC RNA-seq data demonstrate increased levels of alternative splicing in Exp+ FECD. A)** Summary of global differences in splicing events categorised by rMATS. N=4 for controls, N=3 each for FECD Exp+ and FECD Exp-. The most common splice category of alternative splicing observed between all pairwise comparisons was skipped exon, representing ∼70% of all significant alternative splicing events detected. Alt 3’ SS: alternative 3’ splice site, Alt 5’ SS: alternative 5’ splice site, Ret. Intron: retained intron, MXE: mutually exclusive exon **B)** Volcano plot of statistically significant skipped exon events in PWC1 (Control vs Exp+). The dpsi value denotes the magnitude of change for each dysregulated skipped exon event identified. A positive dspi denotes decreased levels of exon inclusion in Exp+, whereas a negative dspi denotes increased levels of exon inclusion in Exp+.

**Figure S3: Pairwise comparison of differentially expressed exons identified by DEXSeq2 demonstrates increased levels of alternative splicing in Exp+ FECD compared to both Exp- FECD and control primary corneal endothelial cells. A)** Table of DEXSeq2 events and genes for each pairwise comparison (PWC). Significance denoted by p-adj ≤ 0.05. **B)** Venn diagram of dysregulated exons coordinates between all three PWCs. The largest overlap is observed between PWC1 and PWC2, highlighting an enrichment of skipped exon events in Exp+ FECD compared to Exp- FECD and controls.

**Table S1:** Clinical details of individuals with Fuchs endothelial corneal dystrophy used to established primary corneal endothelial cell cultures used for downstream analysis.

**Table S2:** Driving genes and additional principal components for PCA analysis of short-read transcriptomic data

**Table S3:** Summary of SQANTI3 analysis of IsoSeq data. In depth characterization of isoforms in control and Exp+ long-read RNA-seq utilising SQANTI3 pipeline following stringent quality control.

**Table S4:** Summary of rMATs results. The top five splice events detected for the following pairwise comparisons (PWC) are presented; (PWC1) control versus Exp+ FECD, (PWC2) Exp+ FECD versus Exp- FECD, and (PWC3) control versus Exp- FECD.

**Table S5:** rMATS identified significant differentially spliced events in Exp+ matching published events with strong association to CTG18.1-expansion mediated FECD.

**Table S6:** Details on the pairwise comparison(s) where rMATS identified new significant differentially spliced events matching published differentially spliced genes with strong association to CTG18.1-expansion mediated FECD.

**Table S7:** Differential gene expression and alternative splicing of fibronectin (*FN1*) in all three pairwise comparisons.

**Table S8:** rMAPS-identified RNAbinding motif enrichment in PWC1 and PWC2, which was also absent in PWC3.

**Table S9:** Differentially expressed genes for all three pairwise comparisons including list of dysregulated genes that are common to FECD in general.

**Table S10:** Pathway enrichment (GO, KEGG, and Reactome) for PWC1, PWC3, and dysregulated genes unique to Exp+.

**Table S11:** Dysregulated RNA binding proteins (RBPs) uniquely in PWC1.

**Table S12:** Summary of all *TCF4* DEXSeq runs

**Table S13:** DEXSeq results for *TCF4* in all three pairwise comparisons showing dysregulation of exons within the gene.

**Table S14:** rMATS *TCF4* mutually exclusive exon events demonstrating a shift between longer isoforms containing 2+ AD domains and shorter isoforms containing 1-2 AD domains detected via alternative splicing pipelines.

**Table S15**: *TCF4* isoform RNAScope sample summary with results of FISH with probe targeting repeat and negative RNAScope control experiments (with probes targeting bacterial genes).

**Table S16:** *TCF4* isoform RNAScope sample summary with results of FISH with probe targeting repeat and negative RNAScope experiments (with probe targeting bacterial genes) in adult human dermal fibroblasts.

**Table S17:** Summary of rare, potentially deleterious, variants in FECD-associated genes identified in Proband B.

**Table S18:** A summary of FRAPOSA-derived ancestry information generated for all CTG18.1 Exp- using genome-wide SNP data extracted from exome sequencing data.

**Table S19:** Clinical data of probands with Fuchs endothelial corneal dystrophy identified to have rare and potentially deleterious heterozygous *TCF4* variants.

## References

1. Mathews P, Benbow A, Corcoran K, DeMatteo J, Philippy B, Van Meter W. 2022 Eye Banking Statistical Report—Executive Summary. Eye Bank Corneal Transplant. 2023 Sep;2(3):e0008.

2. NHSBT. NHS Blood and Transplant Annual Activity Report: Cornea Activity [Internet]. Available from: http://nhsbtdbe.blob.core.windows.net/umbraco-assets-corp/27122/section-10-cornea-activity.pdf

3. Matthaei M, Hribek A, Clahsen T, Bachmann B, Cursiefen C, Jun AS. Fuchs Endothelial Corneal Dystrophy: Clinical, Genetic, Pathophysiologic, and Therapeutic Aspects. Annu Rev Vis Sci. 2019;5(1):151–75.

4. Fautsch MP, Wieben ED, Baratz KH, Bhattacharyya N, Sadan AN, Hafford-Tear NJ, et al. TCF4-mediated Fuchs endothelial corneal dystrophy: Insights into a common trinucleotide repeat-associated disease. Prog Retin Eye Res. 2021 Mar 1;81:100883.

5. Thaung C, Davidson AE. Fuchs endothelial corneal dystrophy: current perspectives on diagnostic pathology and genetics—Bowman Club Lecture. BMJ Open Ophthalmol. 2022 Jul 1;7(1):e001103.

6. Wieben ED, Baratz KH, Aleff RA, Kalari KR, Tang X, Maguire LJ, et al. Gene Expression and Missplicing in the Corneal Endothelium of Patients With a TCF4 Trinucleotide Repeat Expansion Without Fuchs’ Endothelial Corneal Dystrophy. Invest Ophthalmol Vis Sci. 2019 Aug 30;60(10):3636–43.

7. Zarouchlioti C, Sanchez-Pintado B, Hafford Tear NJ, Klein P, Liskova P, Dulla K, et al. Antisense Therapy for a Common Corneal Dystrophy Ameliorates TCF4 Repeat Expansion-Mediated Toxicity. Am J Hum Genet. 2018 Apr 5;102(4):528–39.

8. Depienne C, Mandel JL. 30 years of repeat expansion disorders: What have we learned and what are the remaining challenges? Am J Hum Genet. 2021 May 6;108(5):764–85.

9. Balendra R, Isaacs AM. C9orf72-mediated ALS and FTD: multiple pathways to disease. Nat Rev Neurol. 2018 Sep;14(9):544–58.

10. Wieben ED, Aleff RA, Tang X, Butz ML, Kalari KR, Highsmith EW, et al. Trinucleotide Repeat Expansion in the Transcription Factor 4 (TCF4) Gene Leads to Widespread mRNA Splicing Changes in Fuchs’ Endothelial Corneal Dystrophy. Invest Ophthalmol Vis Sci. 2017 Jan 24;58(1):343–52.

11. Wieben ED, Aleff RA, Tang X, Kalari KR, Maguire LJ, Patel SV, et al. Gene expression in the corneal endothelium of Fuchs endothelial corneal dystrophy patients with and without expansion of a trinucleotide repeat in TCF4. PLOS ONE. 2018 Jul 2;13(7):e0200005.

12. Chu Y, Hu J, Liang H, Kanchwala M, Xing C, Beebe W, et al. Analyzing pre-symptomatic tissue to gain insights into the molecular and mechanistic origins of late-onset degenerative trinucleotide repeat disease. Nucleic Acids Res. 2020 Jul 9;48(12):6740–58.

13. Soragni E, Petrosyan L, Rinkoski TA, Wieben ED, Baratz KH, Fautsch MP, et al. Repeat-Associated Non-ATG (RAN) Translation in Fuchs’ Endothelial Corneal Dystrophy. Invest Ophthalmol Vis Sci. 2018 Apr 1;59(5):1888–96.

14. Westin IM, Viberg A, Byström B, Golovleva I. Lower Fractions of TCF4 Transcripts Spanning over the CTG18.1 Trinucleotide Repeat in Human Corneal Endothelium. Genes. 2021 Dec 17;12(12):2006.

15. Sirp A, Leite K, Tuvikene J, Nurm K, Sepp M, Timmusk T. The Fuchs corneal dystrophy-associated CTG repeat expansion in the TCF4 gene affects transcription from its alternative promoters. Sci Rep. 2020 Oct 28;10(1):18424.

16. Sepp M, Kannike K, Eesmaa A, Urb M, Timmusk T. Functional Diversity of Human Basic Helix-Loop-Helix Transcription Factor TCF4 Isoforms Generated by Alternative 5′ Exon Usage and Splicing. Kashanchi F, editor. PLoS ONE. 2011 Jul 15;6(7):e22138.

17. Zweier C, Peippo MM, Hoyer J, Sousa S, Bottani A, Clayton-Smith J, et al. Haploinsufficiency of TCF4 causes syndromal mental retardation with intermittent hyperventilation (Pitt-Hopkins syndrome). Am J Hum Genet. 2007 May;80(5):994–1001.

18. Amiel J, Rio M, de Pontual L, Redon R, Malan V, Boddaert N, et al. Mutations in TCF4, encoding a class I basic helix-loop-helix transcription factor, are responsible for Pitt-Hopkins syndrome, a severe epileptic encephalopathy associated with autonomic dysfunction. Am J Hum Genet. 2007 May;80(5):988–93.

19. Hellwig M, Lauffer MC, Bockmayr M, Spohn M, Merk DJ, Harrison L, et al. TCF4 (E2-2) harbors tumor suppressive functions in SHH medulloblastoma. Acta Neuropathol (Berl). 2019 Apr 1;137(4):657–73.

20. Stefansson H, Ophoff RA, Steinberg S, Andreassen OA, Cichon S, Rujescu D, et al. Common variants conferring risk of schizophrenia. Nature. 2009 Aug 6;460(7256):744–7.

21. Schizophrenia Psychiatric Genome-Wide Association Study (GWAS) Consortium. Genome-wide association study identifies five new schizophrenia loci. Nat Genet. 2011 Sep 18;43(10):969–76.

22. Zhang D, Dey R, Lee S. Fast and robust ancestry prediction using principal component analysis. Bioinforma Oxf Engl. 2020 Jun 1;36(11):3439–46.

23. Shen S, Park JW, Lu Z xiang, Lin L, Henry MD, Wu YN, et al. rMATS: Robust and flexible detection of differential alternative splicing from replicate RNA-Seq data. Proc Natl Acad Sci. 2014 Dec 23;111(51):E5593–601.

24. Shen S, Park JW, Huang J, Dittmar KA, Lu Z xiang, Zhou Q, et al. MATS: a Bayesian framework for flexible detection of differential alternative splicing from RNA-Seq data. Nucleic Acids Res. 2012 Apr 1;40(8):e61.

25. Park JW, Tokheim C, Shen S, Xing Y. Identifying Differential Alternative Splicing Events from RNA Sequencing Data Using RNASeq-MATS. In: Shomron N, editor. Deep Sequencing Data Analysis [Internet]. Totowa, NJ: Humana Press; 2013 [cited 2022 Oct 31]. p. 171–9. (Methods in Molecular Biology). Available from: 10.1007/978-1-62703-514-9_10

26. Dalton CJ, Lemmon CA. Fibronectin: Molecular Structure, Fibrillar Structure and Mechanochemical Signaling. Cells. 2021 Sep;10(9):2443.

27. Hu J, Rong Z, Gong X, Zhou Z, Sharma VK, Xing C, et al. Oligonucleotides targeting TCF4 triplet repeat expansion inhibit RNA foci and mis-splicing in Fuchs’ dystrophy. Hum Mol Genet. 2018 Mar 15;27(6):1015–26.

28. Konieczny P, Stepniak-Konieczna E, Sobczak K. MBNL proteins and their target RNAs, interaction and splicing regulation. Nucleic Acids Res. 2014;42(17):10873–87.

29. Konieczny P, Stepniak-Konieczna E, Sobczak K. MBNL expression in autoregulatory feedback loops. RNA Biol. 2018 Jan 2;15(1):1–8.

30. Hwang JY, Jung S, Kook TL, Rouchka EC, Bok J, Park JW. rMAPS2: an update of the RNA map analysis and plotting server for alternative splicing regulation. Nucleic Acids Res. 2020 Jul 2;48(W1):W300–6.

31. Love MI, Huber W, Anders S. Moderated estimation of fold change and dispersion for RNA-seq data with DESeq2. Genome Biol. 2014 Dec 5;15(12):550.

32. Mori K, Lammich S, Mackenzie IRA, Forné I, Zilow S, Kretzschmar H, et al. hnRNP A3 binds to GGGGCC repeats and is a constituent of p62-positive/TDP43-negative inclusions in the hippocampus of patients with C9orf72 mutations. Acta Neuropathol (Berl). 2013 Mar;125(3):413–23.

33. Doostparast Torshizi A, Armoskus C, Zhang H, Forrest MP, Zhang S, Souaiaia T, et al. Deconvolution of transcriptional networks identifies TCF4 as a master regulator in schizophrenia. Sci Adv. 2019 Sep 11;5(9):eaau4139.

34. Gerstberger S, Hafner M, Tuschl T. A census of human RNA-binding proteins. Nat Rev Genet. 2014 Dec;15(12):829–45.

35. Li M, Zhuang Y, Batra R, Thomas JD, Li M, Nutter CA, et al. HNRNPA1-induced spliceopathy in a transgenic mouse model of myotonic dystrophy. Proc Natl Acad Sci. 2020 Mar 10;117(10):5472–7.

36. Laurent FX, Sureau A, Klein AF, Trouslard F, Gasnier E, Furling D, et al. New function for the RNA helicase p68/DDX5 as a modifier of MBNL1 activity on expanded CUG repeats. Nucleic Acids Res. 2012 Apr;40(7):3159–71.

37. Pettersson OJ, Aagaard L, Andrejeva D, Thomsen R, Jensen TG, Damgaard CK. DDX6 regulates sequestered nuclear CUG-expanded DMPK-mRNA in dystrophia myotonica type 1. Nucleic Acids Res. 2014 Jun;42(11):7186–200.

38. Anders S, Reyes A, Huber W. Detecting differential usage of exons from RNA-seq data. Genome Res. 2012 Oct;22(10):2008–17.

39. Sznajder ŁJ, Thomas JD, Carrell EM, Reid T, McFarland KN, Cleary JD, et al. Intron retention induced by microsatellite expansions as a disease biomarker. Proc Natl Acad Sci. 2018 Apr 17;115(16):4234–9.

40. Chen W, Wang S, Tithi SS, Ellison DW, Schaid DJ, Wu G. A rare variant analysis framework using public genotype summary counts to prioritize disease-predisposition genes. Nat Commun. 2022 May 11;13(1):2592.

41. ProQR Therapeutics. Open-Label, Single-Dose, Exploratory Study With QR-504a to Evaluate Safety, Tolerability, and Corneal Endothelium Molecular Biomarker(s) in Subjects With Fuchs Endothelial Corneal Dystrophy With Trinucleotide Repeat Expansion in the TCF4 Gene (FECD3) [Internet]. clinicaltrials.gov; 2022 May [cited 2023 Mar 6]. Report No.: NCT05052554. Available from: https://clinicaltrials.gov/ct2/show/NCT05052554

42. Powers A, Rinkoski TA, Cheung K, Schehr H, Osgood N, Livelo C, et al. GeneTAC small molecules reduce toxic nuclear foci and restore normal splicing in corneal endothelial cells derived from patients with Fuchs endothelial corneal dystrophy (FECD) harboring repeat expansions in transcription factor 4 (TCF4). Invest Ophthalmol Vis Sci. 2022 Jun 1;63(7):2753–A0242.

43. Goyer B, Thériault M, Gendron SP, Brunette I, Rochette PJ, Proulx S. Extracellular Matrix and Integrin Expression Profiles in Fuchs Endothelial Corneal Dystrophy Cells and Tissue Model. Tissue Eng Part A. 2018 Apr;24(7–8):607–15.

44. Thériault M, Gendron SP, Brunette I, Rochette PJ, Proulx S. Function-Related Protein Expression in Fuchs Endothelial Corneal Dystrophy Cells and Tissue Models. Am J Pathol. 2018 Jul 1;188(7):1703–12.

45. Angelbello AJ, Benhamou RI, Rzuczek SG, Choudhary S, Tang Z, Chen JL, et al. A Small Molecule that Binds an RNA Repeat Expansion Stimulates Its Decay via the Exosome Complex. Cell Chem Biol. 2021 Jan 21;28(1):34–45.e6.

46. Wang Q, Dou S, Zhang B, Jiang H, Qi X, Duan H, et al. Heterogeneity of human corneal endothelium implicates lncRNA NEAT1 in Fuchs endothelial corneal dystrophy. Mol Ther - Nucleic Acids. 2022 Mar 8;27:880–93.

47. Handsaker R, Kashin S, Reed N, Lee WS, McDonald TM, Tan S, et al. Somatic DNA repeat expansion underlies Huntington’s Disease neuropathology (Plenary Abstract). In Washington DC; 2023.

48. Gall-Duncan T, Sato N, Yuen RKC, Pearson CE. Advancing genomic technologies and clinical awareness accelerates discovery of disease-associated tandem repeat sequences. Genome Res. 2022 Jan;32(1):1–27.

49. Sirp A, Roots K, Nurm K, Tuvikene J, Sepp M, Timmusk T. Functional consequences of TCF4 missense substitutions associated with Pitt-Hopkins syndrome, mild intellectual disability, and schizophrenia. J Biol Chem [Internet]. 2021 Dec 1 [cited 2022 Nov 10];297(6). Available from: https://www.jbc.org/article/S0021-9258(21)01187-X/abstract

50. Tekendo-Ngongang C, Grochowsky A, Solomon BD, Yano ST. Beyond Trinucleotide Repeat Expansion in Fragile X Syndrome: Rare Coding and Noncoding Variants in FMR1 and Associated Phenotypes. Genes. 2021 Oct 22;12(11):1669.

51. Nath S, Caron NS, May L, Gluscencova OB, Kolesar J, Brady L, et al. Functional characterization of variants of unknown significance in a spinocerebellar ataxia patient using an unsupervised machine learning pipeline. Hum Genome Var. 2022 Apr 14;9(1):1–12.

52. Liu F, Liu Q, Lu CX, Cui B, Guo XN, Wang RR, et al. Identification of a novel loss-of-function C9orf72 splice site mutation in a patient with amyotrophic lateral sclerosis. Neurobiol Aging. 2016 Nov 1;47:219.e1–219.e5.

53. Winkler NS, Milone M, Martinez-Thompson JM, Raja H, Aleff RA, Patel SV, et al. Fuchs’ Endothelial Corneal Dystrophy in Patients With Myotonic Dystrophy, Type 1. Invest Ophthalmol Vis Sci. 2018 Jun 1;59(7):3053–7.

54. Mootha VV, Hansen B, Rong Z, Mammen PP, Zhou Z, Xing C, et al. Fuchs’ Endothelial Corneal Dystrophy and RNA Foci in Patients With Myotonic Dystrophy. Invest Ophthalmol Vis Sci. 2017 Sep 1;58(11):4579–85.

55. Wieben ED, Aleff RA, Tosakulwong N, Butz ML, Highsmith WE, Edwards AO, et al. A Common Trinucleotide Repeat Expansion within the Transcription Factor 4 (TCF4, E2-2) Gene Predicts Fuchs Corneal Dystrophy. PLOS ONE. 2012 Nov 21;7(11):e49083.

56. Vasanth S, Eghrari AO, Gapsis BC, Wang J, Haller NF, Stark WJ, et al. Expansion of CTG18.1 Trinucleotide Repeat in TCF4 Is a Potent Driver of Fuchs’ Corneal Dystrophy. Invest Ophthalmol Vis Sci. 2015 Jul;56(8):4531–6.

57. Peh GSL, Chng Z, Ang HP, Cheng TYD, Adnan K, Seah XY, et al. Propagation of human corneal endothelial cells: A novel dual media approach. Cell Transplant. 2015;24(2):287– 304.

58. Babraham Bioinformatics - FastQC A Quality Control tool for High Throughput Sequence Data [Internet]. [cited 2022 Oct 31]. Available from: https://www.bioinformatics.babraham.ac.uk/projects/fastqc/

59. Dobin A, Davis CA, Schlesinger F, Drenkow J, Zaleski C, Jha S, et al. STAR: ultrafast universal RNA-seq aligner. Bioinforma Oxf Engl. 2013 Jan 1;29(1):15–21.

60. Patro R, Duggal G, Love MI, Irizarry RA, Kingsford C. Salmon provides fast and bias-aware quantification of transcript expression. Nat Methods. 2017 Apr;14(4):417–9.

61. Witten DM. Classification and clustering of sequencing data using a Poisson model. Ann Appl Stat. 2011 Dec;5(4):2493–518.

62. Blighe K. PCAtools: everything Principal Component Analysis [Internet]. 2023 [cited 2023 Feb 20]. Available from: https://github.com/kevinblighe/PCAtools

63. Ignatiadis N, Huber W. Covariate powered cross-weighted multiple testing. J R Stat Soc Ser B Stat Methodol. 2021;83(4):720–51.

64. Ignatiadis N, Klaus B, Zaugg JB, Huber W. Data-driven hypothesis weighting increases detection power in genome-scale multiple testing. Nat Methods. 2016 Jul;13(7):577–80.

65. Zhu A, Ibrahim JG, Love MI. Heavy-tailed prior distributions for sequence count data: removing the noise and preserving large differences. Bioinformatics. 2019 Jun;35(12):2084– 92.

66. Raudvere U, Kolberg L, Kuzmin I, Arak T, Adler P, Peterson H, et al. g:Profiler: a web server for functional enrichment analysis and conversions of gene lists (2019 update). Nucleic Acids Res. 2019 Jul 2;47(W1):W191–8.

67. Veiga DT. maseR: Mapping Alternative Splicing Events to pRoteins [Internet]. 2022 [cited 2023 Feb 20]. Available from: https://github.com/DiogoVeiga/maser

68. Gordon SP, Tseng E, Salamov A, Zhang J, Meng X, Zhao Z, et al. Widespread Polycistronic Transcripts in Fungi Revealed by Single-Molecule mRNA Sequencing. PLOS ONE. 2015 Jul 15;10(7):e0132628.

69. Li H. Minimap2: pairwise alignment for nucleotide sequences. Bioinformatics. 2018 Sep 15;34(18):3094–100.

70. Tardaguila M, Fuente L de la, Marti C, Pereira C, Pardo-Palacios FJ, Risco H del, et al. SQANTI: extensive characterization of long-read transcript sequences for quality control in full-length transcriptome identification and quantification. Genome Res. 2018 Jan 3;28(3):396–411.

71. Pontikos N, Yu J, Moghul I, Withington L, Blanco-Kelly F, Vulliamy T, et al. Phenopolis: an open platform for harmonization and analysis of genetic and phenotypic data. Bioinformatics. 2017 Aug 1;33(15):2421–3.

72. McLaren W, Gil L, Hunt SE, Riat HS, Ritchie GRS, Thormann A, et al. The Ensembl Variant Effect Predictor. Genome Biol. 2016 Jun 6;17(1):122.

73. Kircher M, Witten DM, Jain P, O’Roak BJ, Cooper GM, Shendure J. A general framework for estimating the relative pathogenicity of human genetic variants. Nat Genet. 2014 Mar;46(3):310–5.

74. Glusman G, Caballero J, Mauldin DE, Hood L, Roach JC. Kaviar: an accessible system for testing SNV novelty. Bioinformatics. 2011 Nov 15;27(22):3216–7.

75. Jaganathan K, Kyriazopoulou Panagiotopoulou S, McRae JF, Darbandi SF, Knowles D, Li YI, et al. Predicting Splicing from Primary Sequence with Deep Learning. Cell. 2019 Jan 24;176(3):535–548.e24.

76. Zuallaert J, Godin F, Kim M, Soete A, Saeys Y, De Neve W. SpliceRover: interpretable convolutional neural networks for improved splice site prediction. Bioinformatics. 2018 Dec 15;34(24):4180–8.

