## Supplementary material for "Deciphering novel TCF4-driven mechanisms underlying a common triplet repeat expansion-mediated disease": Supplmental information

**Supplemental material**

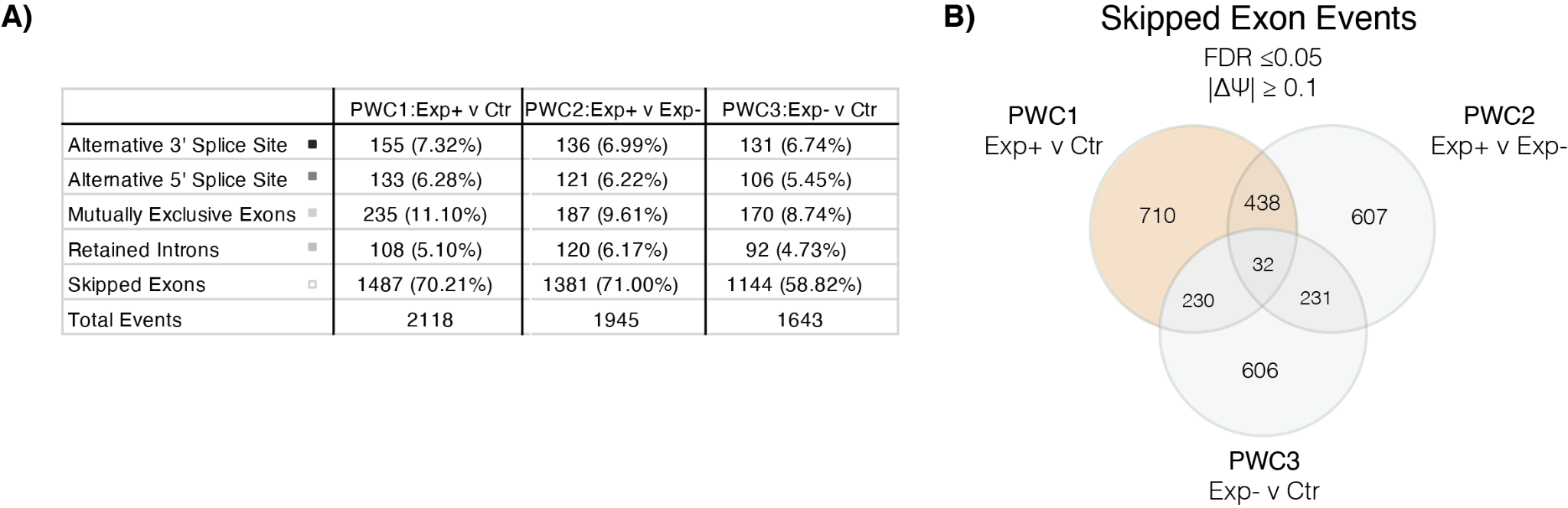

**Figure S1: Pairwise comparison of rMATS alternative splicing analysis demonstrates increased levels of alternative splicing in Exp+ FECD compared to both Exp- FECD and control primary corneal endothelial cells.** Short-read RNA-seq data generated from primary corneal endothelial cells were analysed by rMATS. **A)** Table of rMATS events for each pairwise comparison (PWC). Significance denoted by FDR ≤ 0.05 and deltapsi magnitude larger than 0.1. Values in brackets show percentage of total splice events **B)** Venn diagram of skipped exon event coordinates between all PWCs. The largest overlap is observed between PWC1 and PWC2 highlighting an enrichment of skipped exon events in Exp+ FECD compared to Exp- FECD and controls.

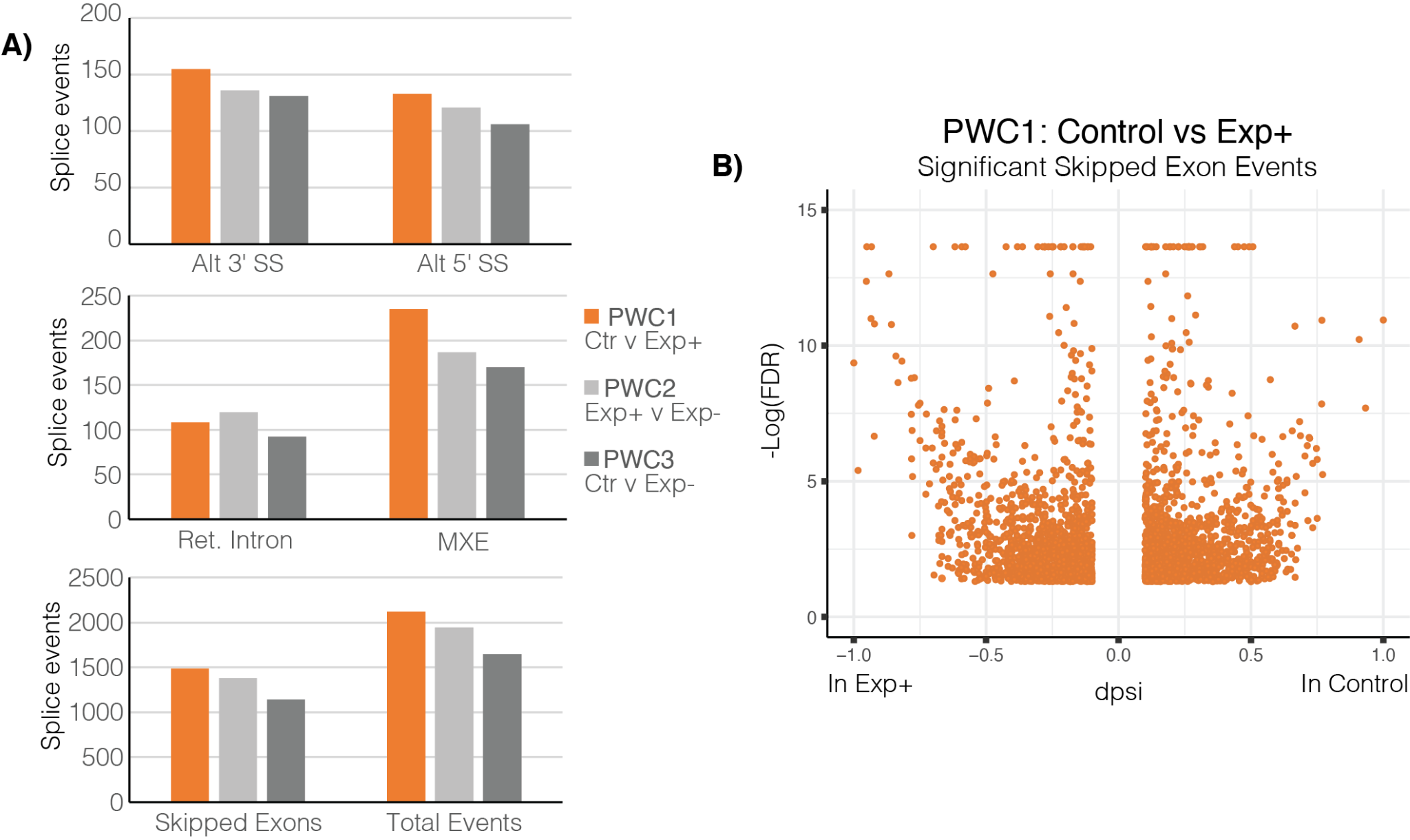

**Figure S2: Alternative splicing analyses of short-read CEC RNA-seq data demonstrate increased levels of alternative splicing in Exp+ FECD. A)** Summary of global differences in splicing events categorised by rMATS**.** N=4 for controls, N=3 each for FECD Exp+ and FECD Exp-. The most common splice category of alternative splicing observed between all pairwise comparisons was skipped exon, representing ~70% of all significant alternative splicing events detected. Alt 3’ SS: alternative 3’ splice site, Alt 5’ SS: alternative 5’ splice site, Ret. Intron: retained intron, MXE: mutually exclusive exon **B)** Volcano plot of statistically significant skipped exon events in PWC1 (Control vs Exp+). The dpsi value denotes the magnitude of change for each dysregulated skipped exon event identified. A positive dspi denotes decreased levels of exon inclusion in Exp+, whereas a negative dspi denotes increased levels of exon inclusion in Exp+.

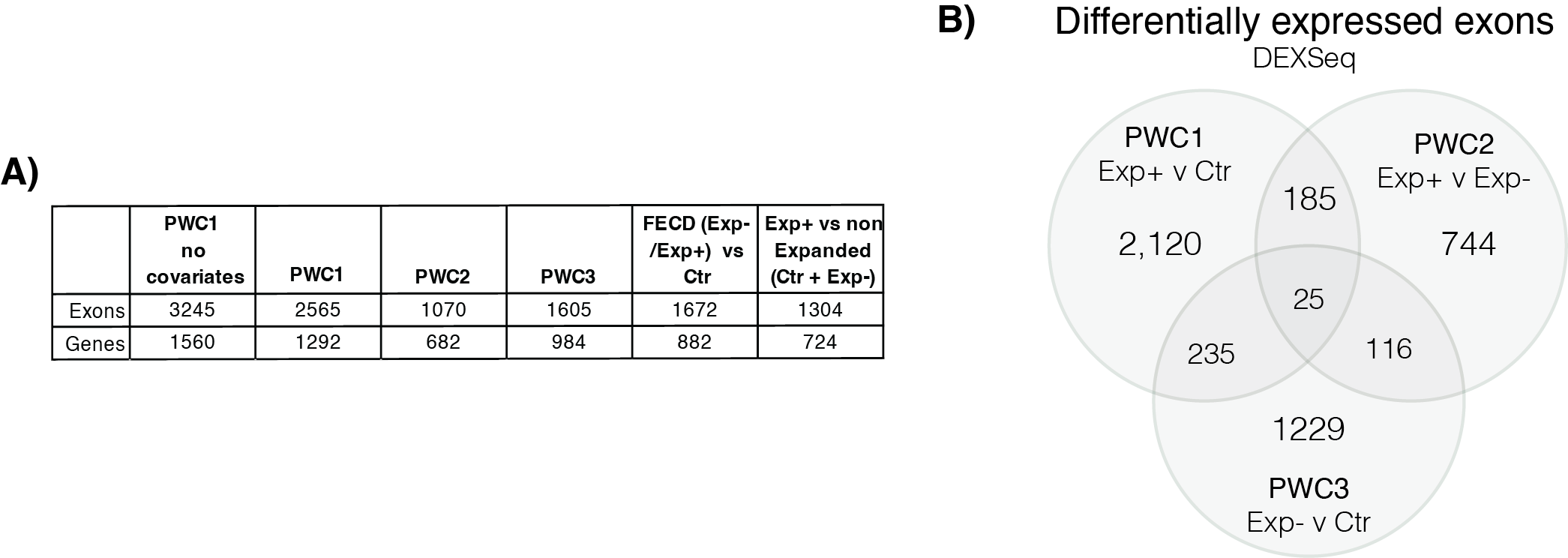

**Figure S3: Pairwise comparison of differentially expressed exons identified by DEXSeq2 demonstrates increased levels of alternative splicing in Exp+ FECD compared to both Exp- FECD and control primary corneal endothelial cells. A)** Table of DEXSeq2 events and genes for each pairwise comparison (PWC). Significance denoted by p-adj ≤ 0.05. **B)** Venn diagram of dysregulated exons coordinates between all three PWCs. The largest overlap is observed between PWC1 and PWC2, highlighting an enrichment of skipped exon events in Exp+ FECD compared to Exp- FECD and controls.

**Table S1: Clinical details of individuals with Fuchs endothelial corneal dystrophy used to established primary corneal endothelial cell cultures used for downstream analysis.**

| **Name/Sex** | **CTG18.1 genotype** | **Age at first keratoplasty (years)*** | **CCT (μm)**  **before surgery** | **Pre-op**  **BCVA** | **CCT (μm)**  **after surgery** | **Final BCVA**** | **Associated ocular features** | **Family history** |
| --- | --- | --- | --- | --- | --- | --- | --- | --- |
| FECD1exp+/F | 43/90 | 72 | NR RE  591 LE | 6/24 RE  6/12 LE | 553 RE  NS LE | 6/9 RE  6/7.6 LE | Cataract BE  ERM peel RE | NK |
| FECD2exp+/M | 14/80 | 78 | 647 RE  640 LE | 6/24 RE  6/36 LE | NS RE  NR LE | 6/7.5 RE  6/9 LE | Cataract BE | NK |
| FECD3exp+/F | 18/86 | 69 | 664 RE  NR LE | 6/9 RE  NR LE | 548 RE  565 LE**** | 6/6 RE  6/9 LE | Cataract BE | NK |
| FECD4exp+/F | 26/78 | 45 | 686 RE  720 LE | 6/5 RE  6/9 LE | NS RE  590 LE | 6/5 RE  6/5 LE | Nil | NK |
| FECD5exp+/F | 16/86 | 67 | 666 RE  681 LE | 6/9 RE  6/18 LE | NR RE  NR LE | 6/6 RE  6/6 LE | Cataract BE | NK |
| FECD6exp+/M | 12/106 | 69 | 620 RE  624 LE | 6/6 RE  6/7.5 LE | 485 RE  489 LE | 6/5 RE  6/4 LE | Cataract BE | 1 sibling |
| FECD7exp+/F | 16/93 | 54 | 633 RE  606 LE | 6/9 RE  7/7.5 LE | 515 RE  508 LE | 6/6 RE  6/5 LE | Cataract BE | NK |
| FECD8exp+/F | 63/88 | 73 | 605 RE  630 LE | 6/18 RE  6/6 LE | 513 RE  NS LE | 6/5 RE  6/6 LE | Cataract BE | NK |
| FECD9exp+/M | 12/76 | 54 | 543 RE  636 LE | RE 6/9  LE 6/12 | NS RE  NR LE | 6/7.5 RE  6/6 LE | Cataract BE | NK |
| FECD10exp+/M | 18/67 | 65 | 587 RE  791 LE | 6/9 RE  6/9 LE | NR RE  NR LE | 6/6 RE  6.7.5 LE | Cataract BE | NK |
| FECD1exp-/F | 13/18 | 61 | 599 RE  588 LE | 6/24 RE  6/9 LE | 690 RE****  NR LE | 6/6 RE  6/9 LE | Cataract BE | NK |
| FECD2exp-/F | 18/24 | 71 | NR RE  699 LE | 6/12 RE  6/12 LE | 7151RE****  572 LE | 6/6 RE  6/5 LE | Cataract BE | NK |
| FECD3exp-/M | 22/28 | 53 | 567 RE  564 LE | 6/12 RE  6/6 LE | 503 RE  NS LE | RE 6/5  LE 6/6 | Cataract BE | NK |
| FECD4exp-/F | 15/15 | 48 | 553 RE  524 LE | 6/18 RE  6/9 LE | 512 RE  498 LE | 6/5 RE  6/5 LE | Cataract BE | NK |
| FECD5exp-/F | 16/23 | 56 | 605 RE  589 LE | 6/12 RE  6/12 LE | 512 RE  504 LE | 6/6 RE  6/6/ LE | Cataract BE | NK |
| FECD6exp-/F | 15/16 | 58 | NR RE  NR LE | 6/9  6/12 | 535 RE  526 LE | 6/7.5 RE  6/7.5 LE | Cataract BE | NK |

M male, F female, CCT central corneal thickness, BCVA best corrected visual acuity, NR no record, NK Not known, NS No surgery performed, RE right eye, LE left eye, BE both eyes, *Descemet membrane endothelial keratoplasty (DMEK) unless stated, **BCVA best corrected visual acuity with spectacle correction (Snellen annotation), **** Descemet stripping automated endothelial keratoplasty (DSAEK), ERM epiretinal membrane

**Table S2: Driving genes and additional principal components for PCA analysis of short-read transcriptomic data**

**(large table)**

**Table S3: Summary of SQANTI3 analysis of IsoSeq data.** In depth characterization of isoforms in control and Exp+ long-read RNA-seq utilising SQANTI3 pipeline following stringent quality control.

| **Characterization of transcripts based on splice junctions** | | | | |
| --- | --- | --- | --- | --- |
|  | Control | | Exp+ FECD | |
|  | # isoforms | # genes | # isoforms | # genes |
| Full Splice Match (FSM) | 1,163 | 1,047 | 3,130 | 2,598 |
| Incomplete Splice Match | 647 | 535 | 2,268 | 1,377 |
| Novel In Catalog (NIC) | 829 | 685 | 1,797 | 1,336 |
| Novel Not In Catalog (NNC) | 437 | 375 | 778 | 639 |
| Genic Genomic | 3 | 3 | 5 | 5 |
| Antisense | 6 | 5 | 4 | 4 |
| Fusion | 24 | 21 | 35 | 33 |
| Intergenic | 4 | 4 | 6 | 6 |
| Terminology based on Tardaguila et al 2018. n= 4 for controls and n=3 for FECD Exp+ (with one sample duplicated to acquire additional depth). FSM: full splice match. ISM: Incomplete splice match. NIC: Novel in catalog. NNC: Novel not in catalog | | | | |

**Table S4: Summary of rMATs results.** The top five splice events detected for the following pairwise comparisons (PWC) are presented; **(PWC1)** control versus Exp+ FECD, **(PWC2)** Exp+ FECD versus Exp- FECD, and **(PWC3)** control versus Exp- FECD.

(large table)

**Table S5. rMATS identified significant differentially spliced events in Exp+ matching published events with strong association to CTG18.1-expansion mediated FECD.**

| **Gene** | **Identified by rMATS (yes/no)** | **rMATS splice type** | **Differential splicing detected via SQANTI3** | **Skipped Exon Coordinates (hg38)** | **dpsi** | **FDR** |
| --- | --- | --- | --- | --- | --- | --- |
| *ABI1^3,6^* | Yes | SE | Yes | chr10:26771075 -26771089 | 0.28 | 1.82E-07 |
| *ADD3^3,6^* | No | - | Yes | - | - | - |
| *AKAP13^3,6^* | Yes | SE | Yes | chr15:85658537 -85658590 | 0.203 | 0.003372 |
| *TSPOAP1^3,6^* | Yes | SE | Excluded | chr17:58308541 -58309380 | 0.222 | 0.028978 |
| *CD46^3,6^* | No | - | Yes | - | - | - |
| *CLASP1^3,6^* | Yes | SE | Yes | chr2:121445449 -121445496 | 0.451 | 0 |
| *COPZ2^3,6^* | Yes | SE | No | chr17:48027676 -48027789 | 0.118 | 7.49E-13 |
| *EXOC1†^3,6^* | Yes | SE | Yes | chr4:55888888 -55888932 | -0.141 | 0.000691 |
| *FGFR1†^3,6^* | Yes | SE | Yes | chr8:38429682 -38429948 | -0.248 | 0 |
| *GOLGA2^3,6^* | Yes | SE | Yes | chr9:128272785 -128272865 | -0.188 | 1.10E-07 |
| *INF2†^3,6^* | Yes | SE | Yes | chr14:104715284 -104715340 | 0.507 | 0 |
| *ITGA6†^3,6^* | Yes | SE | No | chr2:172501772 -172501901 | 0.263 | 0 |
| *KIF13A^a3,6^* | Yes | SE | Excluded | chr6:17789872 -17789910 | 0.17 | 0.00714 |
| *KIF13A^b3,6^* | No | - | Excluded | - | - | - |
| *MBNL1*^3,5,6^* | Yes | SE | Yes | chr3:152446704 -152446757 | -0.15 | 0.000337 |
| *MBNL2*^3,5,6^* | Yes | SE | Yes | chr13:97356796 -97356849 | -0.145 | 3.66E-13 |
| *MYO6^3,6^* | No | - | Yes | - | - | - |
| *NHSL1^3,6^* | Yes | SE | Yes | chr6:138441983 -138442114 | -0.222 | 0.011795 |
| *NUMA1*^3,5,6^* | Yes | SE | Yes | chr11:72012401 -72012442 | 0.268 | 0 |
| *PLEKHM2^3,6^* | Yes | SE | Yes | chr1:15721329 -15721388 | 0.234 | 1.27E-10 |
| *PPFIBP1** | Yes | SE | Yes | chr12:27677064 -27677096 | 0.162 | 7.22E-05 |
| *SCARB1^3,6^* | No | - | Yes | - | - | - |
| *SYNE1*^3,6^* | Yes | SE | Yes | chr6:152145487 -152145555 | 0.223 | 4.36E-09 |
| *VEGFA^3,6^* | No | - | Yes | - | - | - |
| * Events previously validated by RT-PCR, in addition to Mixture of Isoforms (MISO) and MAP-RSeq.^6–8^  † Events where the dysregulated exons match published work, but the flanking exons are not a perfect match.  SE: skipped exon. Samples labelled as excluded from Iso-seq analysis were excluded due to insufficient coverage during sequencing | | | | | | |

**Table S6: Details on the pairwise comparison(s) where rMATS identified new significant differentially spliced events matching published differentially spliced genes with strong association to CTG18.1-expansion mediated FECD.**

| **Gene** | **rMATS splice type** | **Exon Coordinates (hg38)** | **FDR** | **dpsi** | **Verified in Iso-Seq** |
| --- | --- | --- | --- | --- | --- |
| *FGFR1* | MXE | chr8:38429681-38429948, 38457355-38457534 | 0 | 0.125 | Yes |
| *MBNL1* | SE | chr3:1522994040-152300367 | 1.13E-10 | -0.101 | Yes |
| *KIF13A* | SE | chr6:17789871-17789910 | 1.73E-09 | 0.339 | Excluded |
| *KIF13A* | SE | chr6:17794248-17794395 | 1.63E-07 | 0.125 | Excluded |
| *AKAP13* | SE | chr15:85662387-85662453 | 0.0012 | 0.126 | Yes |
| *MBNL2* | SE | chr13:97356795-97356849 | 0.0492 | -0.304 | Yes |
| *MBNL2* | SE | chr13:97366458-97366553 | 0.0011 | -0.114 | Yes |
| *NUMA1* | A5SS | chr11:72068078-72068231 (long), 72068104-72068231 (short) | 0.0441 | 0.113 | No |
| *NUMA1* | SE | chr11:72049418-72049539 | 0.0175 | -0.107 | Yes |
| *TSPOAP1* | SE | chr17:58320108-58320129 | 0.0002 | -0.362 | Excluded |
| MXE: mutually exclusive exon event. SE: skipped exon event. A5SS: alternative 5’ splice site event. Genes excluded from Iso-Seq analysis were done so due to insufficient coverage | | | | | |

**Table S7: Differential gene expression and alternative splicing of fibronectin (*FN1*) in all three pairwise comparisons.**

| **Differential gene expression (DESeq2)** | | | |
| --- | --- | --- | --- |
|  | *shrunkLFC* | *padj* | |
| PWC1 | 2.96419441 | 3.20x10^-20^ | |
| PWC2 | n.s | n.s | |
| PWC3 | 1.9315629 | 3.37x10^-08^ | |
| **Alternative splicing (rMATS)** | | | |
|  | *Dysregulated exons* | *dPSI* | *FDR* |
| PWC1 | Exon 25 (EDB)  Exon 33 (EDA) | -0.253,  -0.305 | 0,  0 |
| PWC2 | Exon 25 (EDB)  Exon 33 (EDA) | 0.262,  0.179 | 0,  4.76E-05 |
| PWC3 | Exon 25 (EDB) | -0.126 | 0.007044931 |
| Exon 25 (EDB) is defined by the genomic coordinates chr2:215,392,931-215,393,203 and Exon 33 (EDA) by chr2:215,380,811-215,381,080. Padj = FDR-adjusted p-value, dPSI=delta percent spliced in, FDR = False discovery rate, n.s. = not significant | | | |

**Table S8: rMAPS-identified RNAbinding motif enrichment in PWC1 and PWC2, which was also absent in PWC3.**

| **GC-containing motifs upregulating skipped exon in Exp+** | |
| --- | --- |
| PCBP2 | CC[CT][CT]CC[ACT} |
| RBM4 | GCGCG[GC][GC] |
| RBM4 | GCGCG[GC]G |
| RBM45 | GACGA[AC][ACG] |
| **TT-containing motifs downregulating skipped exon events in Exp+** | |
| HNRNPC | [ACT]TTTTT[GT] |
| HNRNPCL1 | [ACT]TTTTT[GT] |
| PCBP1 | C[CT]TTCC |
| ZC3H14 | TTT[AGT]TTT |

**Table S9: Differentially expressed genes for all three pairwise comparisons including list of dysregulated genes that are common to FECD in general.**

**(large table)**

**Table S10: Pathway enrichment (GO, KEGG, and Reactome) for PWC1, PWC3, and dysregulated genes unique to Exp+.**

(large table)

**Table S11: Dysregulated RNA binding proteins (RBPs) uniquely in PWC1.**

**Downregulated uniquely in Exp+ (n=35)**

*REXO5,* ***ESRP2****, ZNF106, CPEB3,* ***DDX25****, ENDOU, IFIH1, WARS2, ZNFX1, MRPS7, MIF4GD, RPS4Y1, VARS2,* ***DDX60****, PNPT1, SCAF11, RRNAD1, PARP1, AFF3, PDCD4, TRMT44, MOV10, PPARGC1B, CELF5, ZNF385A, RAVER2, RBM47, DIS3L, CSDC2, NSUN7, PLD6, DDX60L, IFIT1, ELAVL3, NYNRIN*

**Upregulated uniquely in Exp+ (n=148)**

*MSL3, SAMD4A, STAU2, TPR, R3HDM1, RRP12, YBX3, CCAR1, U2AF2, IPO5, AFF4, MBNL3, BZW1, GEMIN5, TUT7, AARS1, LRRFIP2,* ***HNRNPM****, ZC3H14, RBM3, PARP4, GSPT1, POP1,* ***DDX49****, GARS1, EIF3B, UTP6, GAR1, CCDC86, CPSF6, HINT3, RRP9, EIF4G1, PNO1, FARSB, RPF1, PRDX1, HEATR1, PTBP3, ENOX1, UTP20, ZC3H13, SPATS2, XPO5, RIOK1, ATXN1, DGCR8, EIF5A, SYNCRIP, CAPRIN1, IGF2BP3, BZW2, EPRS1, RANBP6, RTCA, SSB, LARP1B, HNRNPA1L2, WARS1, SERBP1, ISG20L2, PNLDC1, NONO, PDCD11,* ***CELF1****, INTS4,* ***MBNL1****, PELO, LARP1, SCAF4, UTP14A, RBPMS, RRP1, ZNF326, DCAF13, NOL6,* ***DDX21****, TRMT61A, URM1, XPO6, RNASEH1, ISG20, MRPL52, AGFG1, CTU2, SRP72, CAVIN1, RRS1, FARSA, NOP10, CNOT10, PCBP3, XPOT, NOC2L, SF3B3, IARS1, YRDC, IPO4, DYNC1H1, TOP1, ARHGEF28, ZNF579, RBM14, EIF6, NA, BOP1*

**Table S12: Summary of all *TCF4* DEXSeq runs**

|  | **PWC1**  **(Exp+ vs Ctr) no covariates** | **PWC1**  **(Exp+ vs Ctr) covariates** | **PWC2**  **(Exp+ vs Exp-)**  **covariates** | **PWC3**  **(Ctr vs Exp-)**  **covariates** | **Exp+ vs non-Expanded**  **(Ctr + Exp-)**  **covariates** | **FECD**  **(Exp-/Exp+ vs Ctr)**  **covariates** |
| --- | --- | --- | --- | --- | --- | --- |
| **Upregulated exons** | E108, E109, E114 | E108, E105 | E013, E014, E015, E083, E084, E108, E109, E110 | n.s | E108, E109 | n.s |
| **Upregulated isoforms** | 6 out of 93 isoforms | 3 out of 93 isoforms | 31 out of 93 isoforms | -- | 4 out of 93 isoforms | -- |
| **Downregulated exons** | E118, E119, E139, E157, E158, | n.s. | E081, E082, E117, E118, E119, E139, E140, E142, E143, E144, E158 | n.s | E119, E157, E158 | n.s |
| **Downregulated isoforms** | 37 out of 93 isoforms | -- | 73 out of 93 isoforms | -- | 63 out of 93 isoforms | -- |
| **Intron retention** | E173, E174 | n.a. |  | n.a. |  | n.a. |

**Table S13: DEXSeq results for *TCF4* in all three pairwise comparisons showing dysregulation of exons within the gene.**

(large table)

**Table S14:** **rMATS *TCF4* mutually exclusive exon events demonstrating a shift between longer isoforms containing 2+ AD domains and shorter isoforms containing 1-2 AD domains detected via alternative splicing pipelines.**

|  | **NM_001083962.2/ENST00000354452.8 exon numbering** | |  |  |
| --- | --- | --- | --- | --- |
| **Splice type event (PWC1)** | **Exon 7** | **Exon 6** | **dpsi** | **FDR** |
| Mutually exclusive exon | chr18:55350873-55351003 | chr18:55403453-55403518 | 0.103 | 0.0023 |
| Mutually exclusive exon | chr18:55350873-55351000 | chr18:55403453-55403518 | 0.102 | 0.0030 |
| **Consequence:** | **Higher in Exp+** | **Higher in Control** |  |  |
|  | **First exon common to both long and short isoforms** | **Last exon common to the majority of long isoforms** |  |  |

**Table S15**: ***TCF4* isoform RNAScope sample summary with results of FISH with probe targeting repeat and negative RNAScope control experiments (with probes targeting bacterial genes).**

| **Sample** | **CTG18.1 repeat genotype** | **CUG**  **specific**  **foci present** | **Negative control clear?** | **Probe B proportion (%)** |
| --- | --- | --- | --- | --- |
| *Unaffected Controls* | | | | |
| Control 8 | 12/16 | No | Yes | 29.43 |
| Control 9 | 15/33 | No | Yes | 28.08 |
| Control 10 | 25/26 | No | Yes | 40.55 |
|  |  |  | Average | 36.02 ± 6.9 |
| *CTG18.1 expansion negative FECD* | | | | |
| FECD4exp- | 15/15 | No | Yes | 27.67 |
| FECD5exp- | 16/23 | No | Yes | 28.18 |
| FECD6exp- | 15/16 | No | Yes | 36.86 |
|  |  |  | Average | 30.90 ± 5.17 |
| *CTG18.1 expansion positive FECD* | | | | |
| FECD8exp+ | 63/88 | Yes | Yes | 15.05 |
| FECD9exp+ | 12/76 | Yes | Yes | 12.58 |
| FECD10exp+ | 18/67 | Yes | Yes | 21.36 |
|  |  |  | Average | 16.33 ± 4.53 |

**Table S16: *TCF4* isoform RNAScope sample summary with results of FISH with probe targeting repeat and negative RNAScope experiments (with probe targeting bacterial genes) in adult human dermal fibroblasts.**

| **Sample** | **Sample info** | **CTG18.1 repeat genotype** | **CUG**  **specific**  **foci present** | **Negative control clear?** | **Probe B proportion (%)** |
| --- | --- | --- | --- | --- | --- |
| *Unaffected Control Adult Human Dermal Fibroblast* | | | | | |
| Control HDF1 | F/62 | 22/27 | No | Yes | 42.83 |
| Control HDF2 | F/46 | 24/27 | No | Yes | 52.37 |
| Control HDF3 | F/38 | 12/23 | No | Yes | 28.30 |
|  | | |  | Average | 41.2 ± 12.1 |
| *CTG18.1 expansion positive FECD human dermal fibroblasts^a^* | | | | | |
| #F1*^a^* | F/69 | 31/69*^a^* | No*^a^* | Yes | 41.81 |
| #F2*^a^* | F/70 | 12/53*^a^* | No*^a^* | Yes | 37.8 |
| #F3*^a^* | F/65 | 84/108*^a^* | No*^a^* | Yes | 38.2 |
| #F4*^a^* | F/78 | 23/68*^a^* | No*^a^* | Yes | 48.3 |
| #F5*^a^* | M/82 | 23/70*^a^* | No*^a^* | Yes | 48.2 |
| #F6*^a^* | M/69 | 12/77*^a^* | No*^a^* | Yes | 50.9 |
|  | | | | Average | 44.2 ± 5.68 |
| *^a^*Data presented from Zarouchlioti et al 2018 | | | | | |

**Table S17:** Summary of rare, potentially deleterious, variants in FECD-associated genes identified in Proband B.

| **Variant (hg38)** | **Gene** | **HGVSc,  HGVSp** | **CADD, MAF gnomAD, MAF Kaviar** | **Proband** |
| --- | --- | --- | --- | --- |
| chr20-3231199-C-T | *SLC4A11* | Het c.1040G>A, p.(Arg347Gln) | 19.24, 1.39 × 10^-3^, 5.2 × 10^-4^ | Proband B 0/1 |
| Exome data from all probands with *TCF4* rare variants were interrogated to determine if rare and potentially deleterious variants in previously reported FECD-associated genes, including *COL8A2*, *SLC4A11*, *AGBL1*, *and ZEB1* could explain their respective disease (MAF < 0.005 in publicly available gnomAD genomes, exomes and Kaviar, CADD score > 15. A single heterozygous *SLC4A11* variant of uncertain significance was identified in Proband B. The annotation from Variant Effect Predictor is filtered to the “most severe” transcripts which is ENST00000380056.3. Abbreviations are as follows: Kaviar, Kaviar Genomic Variant database; gnomAD, The Genome Aggregation Database; UTR, untranslated region; MAF, minor allele frequency; HGVSc, coding DNA sequence based on the Human Genome Variation Society; HGVSp**,** protein sequence based on the Human Genome Variation Society. | | | | |

Table S18:A summary of FRAPOSA-derived ancestry information generated for all CTG18.1 Exp- using genome-wide SNP data extracted from exome sequencing data.

| **Predicted Ethnicity** | N **(%)** |
| --- | --- |
| AFR – African ancestry | 14/134 (10.4%) |
| EAS – East Asian ancestry | 1/134 (0.74%) |
| EUR – European ancestry | 119/134 (88.8%) |

**Table S19: Clinical data of probands with Fuchs endothelial corneal dystrophy identified to have rare and potentially deleterious heterozygous *TCF4* variants.**

| **Age***  **Sex**  **Ethnicity** | **CTG18.1 repeat genotype** | **Age at keratoplasty (years)***** | **CCT (μm)**  **before surgery** | **Pre-op**  **BCVA** | **CCT (μm)**  **after surgery** | **Final BCVA**** | **Associated ocular features** | **Family history (1º relatives)** |
| --- | --- | --- | --- | --- | --- | --- | --- | --- |
| 57/F/black  Proband A | 10/12 | 57 OD  59 OS | 495 OD  493 OS | NA | 441 OD  431 OS  (DMEK) | 0.67 OD  0.67 OS | Cataract surgery | Unknown |
| 42/M/black  Proband B | 17/23 | NP | 589 OD  577 OS | NP | NP | 0.67 OD  0.67 OS | Nil | Unknown |
| 61/F/mixed race (white/black)  Proband C | 13/18 | 61 OD  63 OS | NA  691 OS | NA | 681 OD (DSAEK)  539 OS (DMEK) | 1.0OD  0.67 OS | Cataract surgery | Unknown |
| 74/F/white  Proband D | 12/15 | 75 OD  74 OS | NA | NA | 442 OD  464 OS  (DMEK) | 1.0 OD  1.0 OS | Cataract surgery | Yes |
| 60/F/white  Proband E | 12/16 | 71 OD  60 OS  (PK) | 605 OD NA OS | 0.2 OD 0.25 OS | NA | 0.4 OD 0.6 OS | Cataract surgery,  secondary glaucoma | No |
| 64/F/white Proband F | 12/18 | 73 OD  72 OS | 640 OD  535 OS | 0.5 OD  0.5 OS | 572 OD  527 OS  (DMEK) | 0.67 OD  0.67 OS | Cataract surgery | Yes |
| 56/F/white Proband G | 12/18 | 59 OD  (PK)  63 OS  (DSAEK) 71 OS | NA | NA | 532 OD 539 OS | 0.5 OD 0.8 OS | EBMD, cataract surgery, secondary glaucoma | Yes |

M male, F female, OD right eye, OS left eye, DMEK Descemet membrane endothelial keratoplasty, DSAEK Descemet’s stripping automated endothelial keratoplasty, PK penetrating keratoplasty, CCT central corneal thickness, BCVA best corrected visual acuity, NP corneal surgery not performed, NA not available, EBMD epithelial basement membrane degeneration (Cogan),*Age at diagnosis, **BCVA values converted to decimal annotation, ***DMEK unless stated.
